# Midbrain Dopaminergic Neuron Development is Regulated by Two Molecularly Distinct Subtypes of Radial Glia Cells

**DOI:** 10.1101/2024.03.08.584031

**Authors:** Emilía Sif Ásgrímsdóttir, Luca Fusar Bassini, Pia Rivetti di Val Cervo, Daniel Gyllborg, Clàudia Puigsasllosas Pastor, Kawai Lee, Christopher Grigsby, Saiful Islam, Peter Lönneberg, Carlos Villaescusa, Carmen Saltó, Sten Linnarsson, Gioele La Manno, Enrique M. Toledo, Ernest Arenas

## Abstract

Understanding midbrain dopaminergic (mDA) neuron development is key to advancing cell replacement therapies for Parkinson’s disease (PD). Recent single-cell RNA-sequencing (scRNA-seq) studies have identified different subtypes of transient radial glia (Rgl) cell types in the developing mouse and human ventral midbrain. However, their individual functions and their impact on mDA neuron development are unclear. Here we analyze the transcriptome of endogenous mouse and human ventral midbrain Rgl and assess the function of key midbrain floor plate Rgl factors in human stem cells during mDA neuron differentiation. We find that Rgl1 is defined by a neurogenic network centered on *Arntl*, and that this transcription factor regulates human mDA neurogenesis. Conversely, the transcriptome of Rgl3 is dominated by signaling and extracellular matrix molecules that control different aspects of human mDA neuron development. Thus, our results suggest a function of Rgl1 as a mDA progenitor, and of Rgl3 as a signaling mDA niche cell. Moreover, using human stem cells we demonstrate that new knowledge of cell-type specific intrinsic and extrinsic developmental factors can readily be applied to improve the generation of cells with therapeutic interest, such as human mDA neurons for PD.

## Introduction

Midbrain dopaminergic (mDA) neurons are the primary cells affected by Parkinson’s disease (PD) and understanding their development has been crucial to advancing PD cell replacement therapy (Arenas et al., 2015; Garritsen et al., 2023). mDA neurons are born amongst radial glia (Rgl) in the floor plate (FP) region of the ventral midbrain (VM), and their development is controlled by the interplay of two signaling centers, the isthmic organizer at the midbrain-hindbrain boundary (Joyner et al., 2000), and the midbrain floor plate (mFP) itself (Placzek and Briscoe, 2005). In recent years, single-cell RNA-sequencing (scRNA-seq) technologies have revealed the cellular diversity of the developing VM and provided novel insights into the molecular programs that regulate mDA neuron development (Braun et al., 2023; Kee et al., 2017; La Manno et al., 2021, 2016; Tiklová et al., 2019; Xie et al., 2023). However, while our understanding of cell types in the VM has grown considerably, the specific functions of signaling factors and extracellular matrix (ECM) derived from individual cell types on mDA neuron development and the components that make up the cellular microenvironment or dopaminergic niche remain to be elucidated.

Found throughout the entire developing CNS, Rgl cells constitute a heterogeneous group of transient cells that contribute to different aspects of CNS development (Barry et al., 2014; Campbell and Götz, 2002; Kriegstein and Götz, 2003). First observed by Camilo Golgi with silver impregnations, Rgl cells were suggested by Giuseppe Magini to be a glial scaffold for neurons migrating from the ventricular neuroepithelium towards the mantle zone (Reviewed in Bentivoglio and Mazzarello, 1999), a function that was later confirmed by electron microscopy studies (Rakic, 1981). Rgl cells were subsequently recognized to be neuronal progenitors (Rakic, 1988), and are currently considered as neural stem cells capable of limited self-renewal and of giving rise to both neurons and glia (Malatesta and Götz, 2013). However, the function of Rgl varies along the dorsoventral and anterior-posterior axis of the developing CNS. In the FP, Rgl are primarily considered as non-neurogenic cells capable of secreting factors that direct neuronal identities and axonal trajectories (Placzek and Briscoe, 2005). However, mFP Rgl differ from all other FP cells along the neuraxis in that they also express roof plate genes (Arenas et al., 2015; Garritsen et al., 2023), and can undergo neurogenic divisions to give rise to mDA neurons (Bonilla et al., 2008; Ono et al., 2007). More recently, scRNA-seq analysis of the developing mouse and human VM identified three main molecular subtypes of Rgl, Rgl1-3, with unique gene expression profiles and spatial distributions (Braun et al., 2023; La Manno et al., 2016). However, it is unclear whether their functions in the developing midbrain are identical or distinct, and how they influence DA neuron development.

To address these fundamental questions and gain a further understanding of the cell types as well as the cell-intrinsic and extrinsic factors that regulate mDA development, we analyzed single-cell and bulk RNA-seq data from the developing mouse and human VM. Our study identifies Rgl1 and Rgl3 as the two main Rgl cells contributing to the mDA niche, with each cell type having distinct functions. We identify Rgl1 as a neurogenic cell type, and Rgl3 as the main contributor of cell-extrinsic factors, including growth factors, morphogens, and ECM. We use human embryonic and neuroepithelial stem cells to experimentally demonstrate that cell-intrinsic factors expressed by Rgl1 regulate mDA neurogenesis whereas cell-extrinsic factors expressed by Rgl3 regulate diverse aspects of mDA neuron development. These findings were further validated through computational analysis of endogenous human VM datasets. Our study thus represents the first unbiased analysis of the contribution of mouse and human Rgl subtypes to mDA neuron development, identifies highly specialized functions of Rgl subtypes, as well as cell-intrinsic and -extrinsic factors that may find a future application in cell replacement therapies for PD.

## Results

### Transcriptomic analysis of the embryonic ventral midbrain: defining the mDA niche

To identify gene-expression networks that define the mDA niche and distinguish the developing VM from adjacent brain compartments, we performed bulk RNA-seq with embryonic mouse VM, ventral hindbrain (HB), ventral forebrain (FB), dorsal midbrain (DM) and alar plate (L), from day E11.5 to E14.5 (Fig. 1A). Specific anatomical and temporal gene expression patterns were defined (Fig. S1A) and multiple differentially expressed genes (DEG) were identified in the VM compared to other brain regions (Table S1). A pair-wise correlation of all VM samples revealed E11.5 as the most divergent stage (Fig. S1B), and principal component analysis (PCA) confirmed the similarity between E12.5 and E13.5 (Fig. S1C). Unbiased weighted gene co-expression network analysis allowed us to identify a molecular signature that defined and differentiated the developing VM from neighboring brain regions. One of the 13 modules of co-expressed genes (light-green) was enriched in most VM DEGs from E11.5 to E14.5 (Fig. S2A and Table 1). Moreover, the top 5% of VM interactions, by strength of the correlation (1235 out of 24506), involved 48% of genes in this module (181 out of 374), which were enough to separate the VM samples from neighboring brain regions at all time points analyzed, as assessed by PCA (Fig. S2B). This refined module contained many genes expressed in the mDA lineage (*Foxa1*, *Shh*, *Wnt5a*, *Th*, *Slc6a3*, *Pitx3*, etc.) and was thus designated as the mDA module (Fig. S2C). Gene co-expression network analysis of the mDA module identified correlated genes (edge-connected nodes) with similar expression patterns and changes over time (color) (Fig. 1B). GO term analysis of this network identified several biological processes relevant to midbrain development (Fig. S2D), such as dopamine biosynthetic process, neural tube development, and neuron projection guidance. Notably, ECM was amongst the most significantly enriched GO terms, suggesting a prominent function in mDA neuron development.

**Figure 1.**
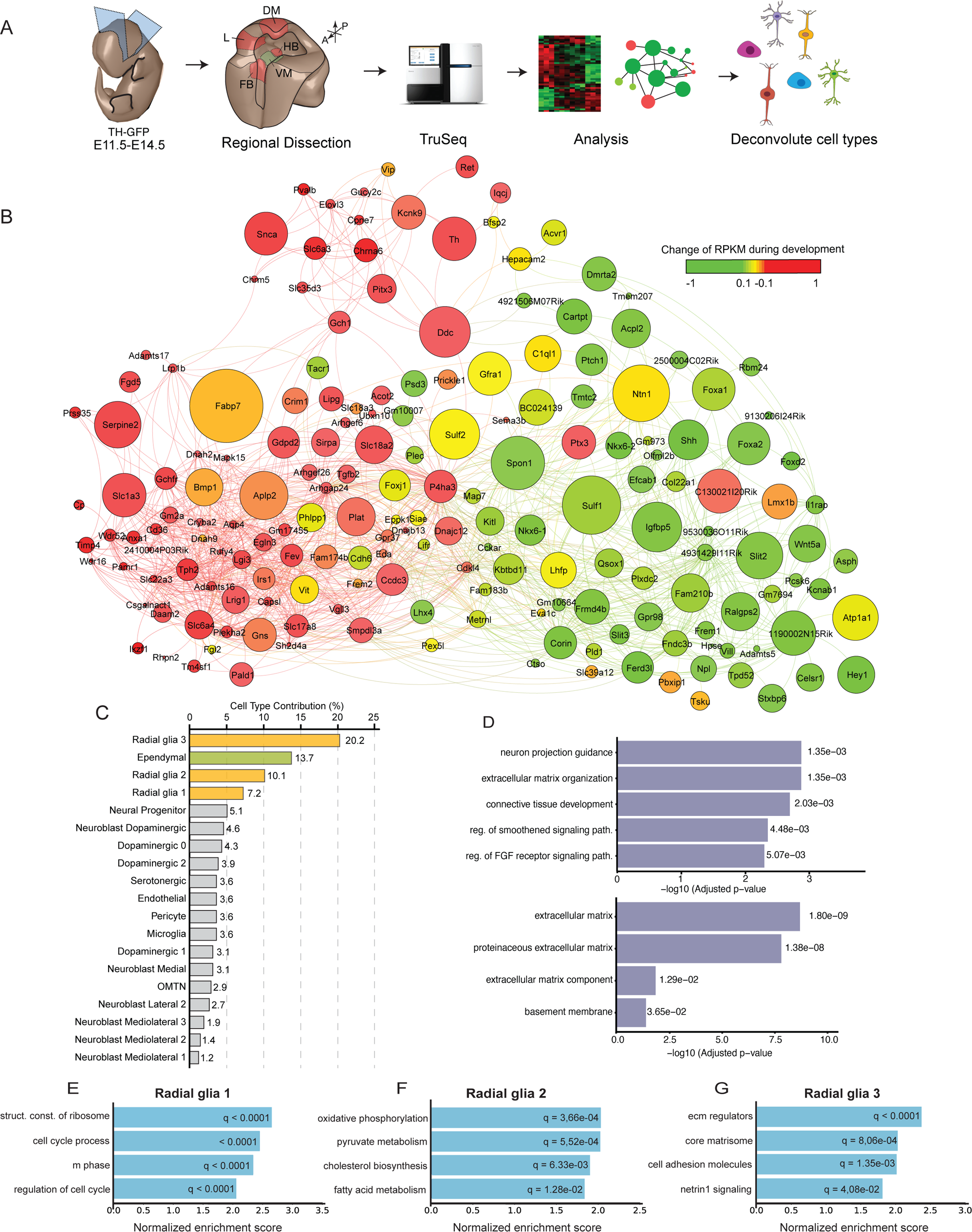
Transcriptomic profile of the ventral midbrain: the dopaminergic module. **(A)** Schematic overview of workflow. Embryonic tissue was collected for RNA-seq from TH-GFP mouse embryos of stages E11.5, E12.5, E13.5 and E14.5. VM, ventral midbrain. DM, dorsal midbrain. FB, forebrain floor plate. HB, hindbrain floor plate. L, alar plate. Bulk RNA-seq data was used to define the transcriptional network of the VM (dopaminergic module) and scRNA-seq data from the developing VM was used to deconvolute the dopaminergic module and assign cell types to the expression profile/s in the module. **(B)** Weighted gene co-expression network analysis of the mDA module, filtered for the top 5% of interactions. Color represents changes in levels of gene expression during development. Node size is proportional to the mean expression levels of the gene during development. **(C)** Contribution of individual VM cell types to the transcriptional network formed by the dopaminergic module. **(D)** GO terms corresponding to the transcriptional profile of the top four cell types contributing to the mDA module (Rgl1-3 and ependymal cells). Top, GO for biological process. Bottom, GO for cellular components. **(E,F,G)** Gene set enrichment analysis (GSEA) of Rgl1, 2 and, 3 transcriptomes. Selected terms corresponding to top normalized enrichment scores (NES) shown for Rgl1 **(E)**, Rgl2 **(F)**, and Rgl3 **(G)**.

**Table 1.**

| Module | Number<br>VM DEG | Module<br>size | q-value | Enrichmen<br>t Score |
| --- | --- | --- | --- | --- |
| Black | 37 | 551 | 9.12e-02 | -0.55 |
| Brown | 46 | 856 | 1.08e-01 | -0.14 |
| Cyan | 6 | 281 | 9.15e-04 | -5.68 |
| Green | 119 | 2839 | 9.26e-03 | 6.10 |
| Green Yellow | 10 | 423 | 1.80e-04 | -6.35 |
| Grey | 24 | 519 | 3.99e-02 | -1.53 |
| Grey60 | 55 | 175 | 1.03e-24 | 5.80 |
| Light cyan | 18 | 201 | 3.26e-02 | -2.01 |
| Light green | 191 | 378 | 5.88e-136 | 824.97 |
| Light yellow | 2 | 100 | 2.46e-02 | -3.28 |
| Magenta | 69 | 3087 | 3.56e-17 | 13.90 |
| Salmon | 54 | 287 | 6.10e-14 | 2.63 |
| Tan | 11 | 371 | 2.63e-03 | -4.27 |

The mDA module was deconvoluted using scRNA-seq analysis of the developing VM (La Manno et al., 2016) by assigning cell types to the expression profiles in the mDA network. 97% of the genes in the mDA network (176 of 181 genes) could be assigned to at least one midbrain cell type (Fig. S3). Surprisingly, 51.2% of the genes in the mDA module were contributed by Rgl1-3 and ependymal cells (Fig. 1C), suggesting a major role for these cells in the mDA niche. GO term analysis of these four cell types underlined their contribution to ECM components, neuron projection guidance, as well as Shh and Fgf signaling (Fig. 1D).

Since Rgl1-3 are selectively found in the VM (La Manno et al., 2016), we performed gene set enrichment analysis (GSEA) of these cells. Rgl1 was enriched in genes related to proliferation (Fig. 1E), Rgl2 in mitochondrial respiration and lipid metabolism (Fig. 1F), and Rgl3 in ECM components as well as signaling pathways (Fig. 1G). These results suggested a functional specialization of Rgl in the mDA niche and led us to examine their individual contributions in further detail.

### Identification of a transcriptional network in Rgl1 involved in mDA neurogenesis

To further characterize the role of each Rgl, we curated a database of transcription factors (TFs) expressed by Rgl1-3 and their known target genes (Janky et al., 2014). We then clustered the TFs and target genes expressed by each Rgl (Fig. 2A and Fig. S4A-B) and investigated which TF combinations can most effectively explain their transcriptional profiles. TFs with upregulated target genes in each Rgl combined with a Fast Westfall-Young (FWY) permutation test for multiple testing correction (Terada et al., 2013a, 2013b) were used to identify statistically significant TF combination patterns (Fig. 2B). Notably, the FWY procedure prunes for non-significant combinations (Tarone, 1990), transforming the combinatorial analysis of downstream target genes into a tractable computational problem. As a control we performed FWY analysis on TFs randomly selected from the molecular signature database repository (MSigDB) while maintaining the transcriptional profile of each Rgl. This resulted in much fewer statistically significant combinations than obtained using the TFs defining each Rgl (Fig. S4C), validating the specificity of our analysis.

**Figure 2.**
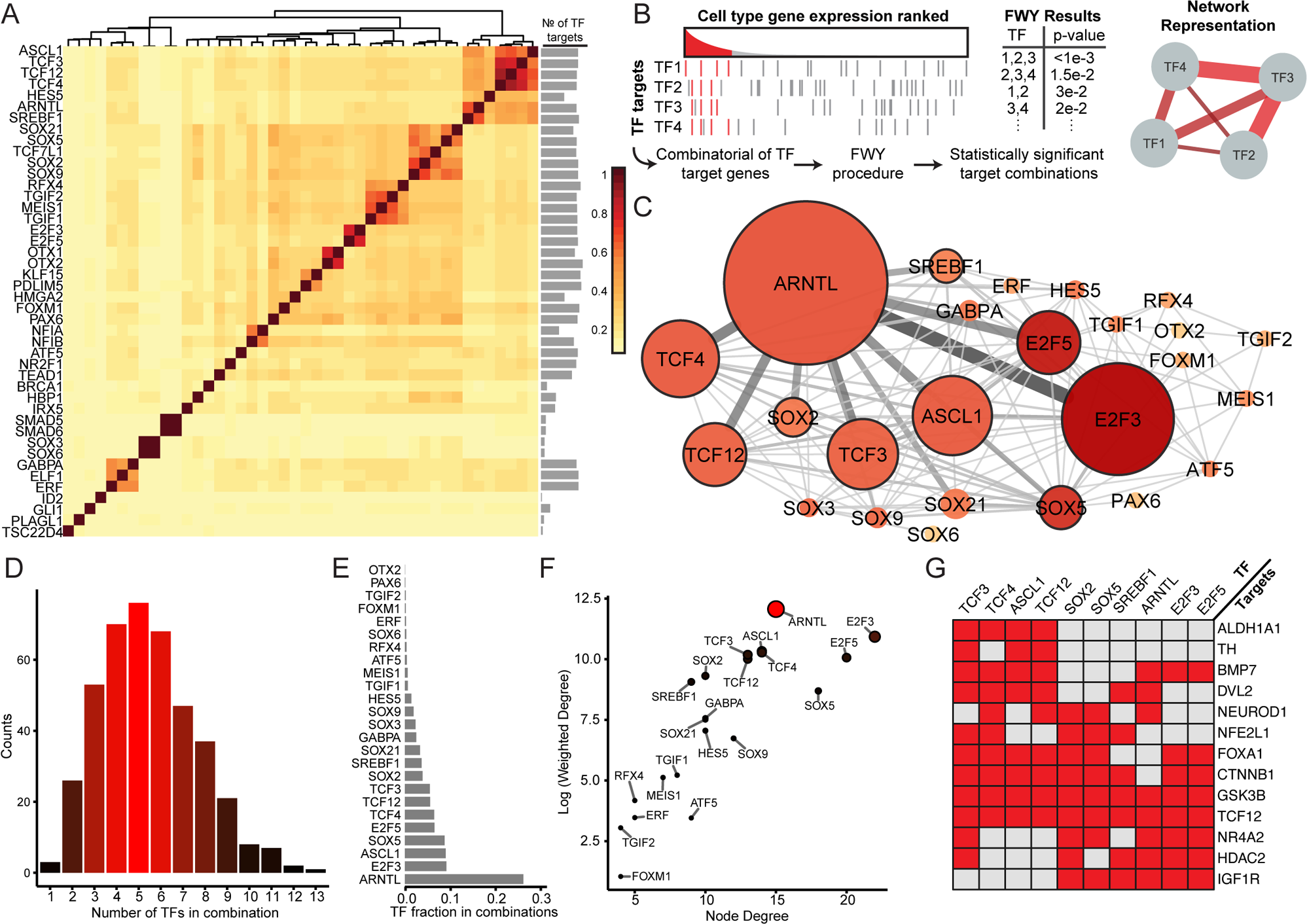
Combinatorial analysis of transcription factors defines the transcriptional network of Rgl1. **(A)** Clustering of transcription factors expressed in Rgl1 by Jaccard index of shared target genes. A bar plot to the right represents the number of targets genes for each transcription factor. **(B)** Diagram illustrating the combinatorial analysis of transcription factor (TF) target genes used to define the transcriptional network. Left, enriched target genes are defined as genes regulated by TF1-4 and expressed above threshold. Red lines represent upregulated target genes. Several target genes were shared by different TFs. Right, network illustrating TFs with shared upregulated target genes analyzed by Fast Westfall-Young (FWY) multiple combinatorial transcription factor enrichment. Transcription factors that regulate the same or overlapping sets of target genes are connected in the network. **(C)** Rgl1 transcriptional network generated by FWY analysis. Node color reflects the number of transcription factor connections (node degree) and node size is proportional to the strength of the connection (weighted degree). Nodes with high weighted degrees (core nodes) have a black border. Color intensity and width of the line connecting two nodes (edge) is proportional to the interaction score of the TF pair. **(D)** Distribution of the number of TFs involved in significant combinations. **(E)** Distribution of significant TFs based on the frequency of detection for a particular TF in a TF combination. **(F)** Plot comparing node degree and weighted node degree of the network. For presentation purpose, weighted degree is shown in logarithmic scale. Color intensity and size is proportional to weighted degree. **(G)** Heat map of selected target genes regulated by core transcription factors in the Rgl1 TF network. Target genes defined by the database by Janky et al. 2014.

Analysis of Rgl1, the earliest appearing Rgl in the developing VM (La Manno et al., 2016) defined two large Sox and pro-neural basic helix-loop-helix (bHLH) TF clusters with common target genes (Fig. 2A), and FWY analysis revealed that 25 TFs generate 419 significant combinations (Table S2). We then constructed a TF network by calculating a pairwise interaction score for all significant TF combinations, based on frequency and p-value (Fig. 2C). The mean number of TFs per combination was 5.465 (Fig. 2D) and the five most common TFs were *Arntl/Bmal1*, *E2f3*, *Ascl1*, *Sox5* and *E2f5* (Fig. 2E). *E2f5* and *E2f3,* two cell cycle regulators (Chen et al., 2009; Trimarchi and Lees, 2002), had the highest number of interaction partners (node degree), but these interactions were less frequent and with higher p-values. When node degrees were weighted for interaction score, *Arntl* was identified as the most relevant gene in the network (Fig. 2F), followed by *E2f3, Ascl1, Sox5*, *E2f5, Tcf4*, *Tcf12*, *Tcf3, Sox2* and *Srebf1*, indicating that components of the bHLH neurogenic cluster explain most aspects of the transcriptional state of Rgl1. Moreover, several genes in this cluster have previously been found to regulate mDA neurogenesis, including *Ascl1,* which works in concert with *Neurog2* (Kele et al., 2006)*; Srebf1,* which is both required and sufficient for mDA neurogenesis (Toledo et al., 2020); and *Tcf3*/*4*, that interact with active β-catenin to control mDA neurogenesis (Arenas, 2014; Nouri et al., 2020).

The downstream target genes of this neurogenic network were also analyzed using curated gene sets from MSigDB (Subramanian et al., 2005). Cell cycle regulation, as well as Wnt and Notch signaling were found to be the main functions controlled by the network (Fig. S4D), all of which are fundamental for mDA neurogenesis (Arenas et al., 2015; Louvi and Artavanis-Tsakonas, 2006). Putative target genes known to regulate mDA neuron development included components of the Wnt pathway, such as *Dvl2*, *Ctnnb1*, *Gsk3b* and *Tcf12* (Inestrosa and Arenas, 2010; Neuman et al., 1993; Willert and Jones, 2006), and of other pathways, such as *Foxa1* and *Bmp7* (Hynes et al., 1995; Jovanovic et al., 2018), and *Igf1r* (Quesada et al., 2007) (Fig. 2G). Furthermore, we identified genes involved in neurogenesis, such as *Neurod1* (Cho and Tsai, 2004; Dennis et al., 2019), *Hdac2* (Jawerka et al., 2010; Montgomery et al., 2009), and *Tcf12* (Mesman and Smidt, 2017; Uittenbogaard and Chiaramello, 2002), as well as TFs required for mDA neuron development, such as *Nfe2l1* (Villaescusa et al., 2016) and *Nr4a2* (Zetterström et al., 1997) and markers of mature mDA neurons, including *Th* and *Aldh1a1* (Arenas et al., 2015; Galter et al., 2003). Thus, our results indicate that a transcriptional network mainly formed by *Arntl, Ascl1*, *Tcfs* (*3*, *4* and *12*), *Soxs* (*2* and *5*) and *Srebf1* can set in motion a transcriptional program that controls cell cycle exit and neurogenesis, and allows Rgl1 to respond to Wnt and Shh signaling and differentiate into postmitotic mDA neurons.

### ARNTL, the central node of the transcription factor network in Rgl1, controls mDA neurogenesis

Our analysis identified *Arntl* as the central and most predominant TF in the network that explains the transcriptional profile of Rgl1. *Arntl* is known to promote neurogenesis, control cell cycle exit, and regulate circadian rhythm (Bouchard-Cannon et al., 2013; Kimiwada et al., 2009; Malik et al., 2015). However, it is unknown whether *Arntl* plays any role in VM development or mDA neurogenesis, as suggested by our data. We started by examining Arntl protein in the developing VM and found it in FP Sox2^+^ cells at E13.5 (Fig. S5A). To investigate a potential function of *Arntl* in mDA neurogenesis, we utilized a human long-term neuroepithelial stem (hLT-NES) cell line (Sai2, Tailor et al., 2013), for gain and loss of function experiments during mDA neuron differentiation (Fig. 3A). As previously reported, hLT-NES cells acquired a mFP phenotype (Villaescusa et al., 2016) as assessed by the presence of TH^+^ neurons co-expressing the mDA markers FOXA2, LMX1A and NR4A2 at day 8 of differentiation (Fig. 3B-D).

**Figure 3.**
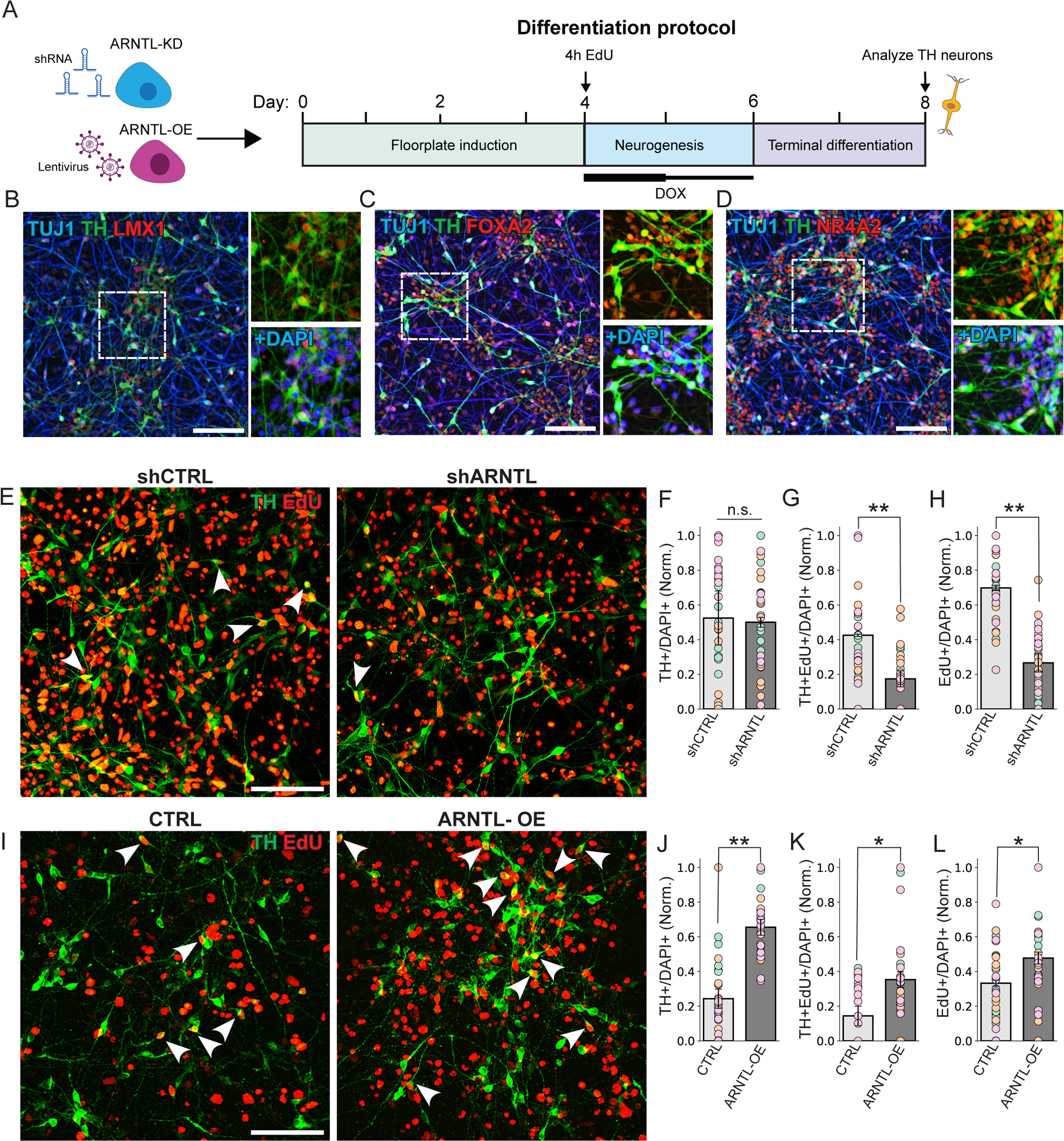
ARNTL influences the dynamics of mDA neurogenesis in human neuroepithelial stem cells. **(A)** Schematic representation of the differentiation protocol used to study dopaminergic neurogenesis in stable ARNTL knockdown (KD) and overexpression (OE) hLT-NES cells. ARNTL overexpression was induced by doxycycline on day 4 (2 µg/ml) and day 5 (1 µg/ml). (**B, C, D**) Immunostaining of LMX1A^+^; TH^+^ (**B**), FOXA2^+^; TH^+^ (**C**), and NR4A2^+^; TH^+^ (**D**) neurons on day 8 confirms the accurate patterning of mDA neurons derived from hLT-NES. TUJ1 is included for visualization of neuronal projections. Scale bar, 100 µm. **(E)** DA neurogenesis was analyzed in shControl or shARNTL hLT-NES by identifying Th^+^EdU^+^ (arrows) neurons. Cells were pulsed with EdU for 4 hours on day 4, and EdU detection was performed on day 8. Scale bar, 100 µm. **(F, G and H)** Quantification of TH^+^/DAPI^+^, (shARNTL = 0.52 ± 0.03, mean ± SEM; shCTRL = 0.5 ± 0.16) **(F),** TH^+^ EdU^+^/DAPI ^+^, (shCTRL = 0.69 ± 0.02; shARNTL = 0.27 ± 0.05, p = 0.0057) **(G)** and EdU^+^/DAPI^+^ (shCTRL = 0.42 ± 0.006; shARNTL = 0.17 ± 0.05, p = 0.0015) **(H)** on day 8, normalized to the average of each experiment. *, p < 0.05, **p < 0.01, student’s t-test. Each data point is represented by a circle, and the circle’s color corresponds to the biological replicate (n = 3 independent experiments). (**I-L**) Analysis of DA neurogenesis in hLT-NES overexpressing EGFP (CTRL) or ARNTL (ARNTL-OE). ARNTL or EGFP overexpression was induced with doxycycline (2 µg/ml) on day 4 and maintained on day 5 (1 µg/ml). Cells were pulsed with EdU for 4 hours on day 4. (**I)** EdU detection and TH immunocytochemistry was performed on day 8. Scale bar, 100 µm. **(J, K and L)** Quantification of TH^+^/DAPI^+^ (CTRL = 0.24 ± 0.05; ARNTL-OE = 0.65 ± 0.04, p = 0.0046) (**J**), TH^+^EdU^+^/DAPI^+^ (CTRL = 0.14 ± 0.05; ARNTL-OE = 0.35 ± 0.05, p =0.043) (**K**), and EdU^+^/DAPI^+^ (CTRL=0.33 ± 0.02 mean ± SEM; ARNTL-OE=0.48 ± 0.03, p = 0.02) (**L**) on day 8, normalized to the average of each experiment. *, p < 0.05, **p < 0.01, student’s t-test. Each data point is represented by a circle, and the circle’s color corresponds to the biological replicate (n = 3 independent experiments).

A stable *ARNTL* knock-down (KD) cell line (shARNTL) exhibiting dramatically reduced levels of ARNTL compared to a control shRNA cell line (shCTRL, Fig. S5B) was generated. shARNTL cells were pulsed with EdU for 4h at day 4 of mDA differentiation to label proliferating cells and examine neurogenesis (Fig. 3A). While we did not observe a change in the proportion of TH^+^ neurons in response to ARNTL KD (Fig. 3E,F), the proportion of cells undergoing proliferation (EdU^+^/DAPI^+^) or mDA neurogenesis (TH^+^EdU^+^/DAPI^+^) decreased by 61% and 60%, respectively (Fig. 3E,G,H). Notably, analysis of cells before the EdU pulse, at day 4, revealed fewer proliferative Ki67^+^ cells and more TH^+^ neurons in the shARNTL condition compared to the control (Fig. S5C). These results indicate that *ARNTL* KD does not affect the total number of mDA neurons but rather the timing of cell cycle exit and mDA neurogenesis, occurring prior to day 4 in ARNTL KD and during day 4-8 in control cells.

To confirm the function of ARNTL, we generated stable, tetracycline-inducible ARNTL or EGFP hLT-NES cell lines (Fig. S5D). Cells were differentiated towards mDA neurons, treated with doxycycline (2 and 1 µg/mL at days 4 and 5, respectively), pulsed with EdU (4h at day 4), and examined at day 8 (Fig. 3A). We found that ARNTL expression increased proliferating progenitor cells by 45% (EdU^+^/DAPI^+^), the proportion of cells undergoing mDA neurogenesis from day 4-8 by 150% (EdU^+^TH^+^/DAPI^+^), and the proportion of TH^+^ neurons by 171% compared to control (Fig. 3I-L). Combined, our results show that *ARNTL,* the central node in the Rgl1 transcriptional network, regulates both the timing of progenitor cell cycle exit and mDA neurogenesis, with sh*ARNTL* leading to reduced proliferation and early mDA neurogenesis, while *ARNTL*-OE increased progenitor proliferation and subsequent mDA neurogenesis.

### Transcriptional networks in Rgl2 and 3

Analysis of TFs enriched in Rgl2 identified two Sox and pro-neural bHLH TF clusters (Fig. S4A). However, when investigating the TF combinations that best explain the transcriptional profile of Rgl2, FWY revealed that the only significant combination was *Pax6* and *Tcf7l1* (p-value=0.025), a combination that has been reported to maintain neural progenitors in an undifferentiated state (Kuwahara et al., 2014; Thakurela et al., 2016). Analysis of the target genes for these two TFs reveled enrichment of GO terms associated to development of the forebrain (p-value: 3.12e-45), hindbrain (p-value: 7.88e-8) and midbrain (p-value: 5.89e-5), suggesting a generic role of this network in neural progenitor maintenance. These findings together with the fact that Rgl2 cells and *Pax6* expression are found lateral to the mFP, in the basal plate of the midbrain (La Manno et al., 2016), suggests that the function of Rgl2 is unlikely to be directly related to mDA neuron development.

Conversely, Rgl3 is found in the mFP (La Manno et al., 2016), and our analysis identified two different clusters with Sox and bHLH genes (Fig. S4B). FWY analysis of Rgl3 revealed 15 enriched TF combinations (Table S2), forming a network centered on *Tead1*, a component of the hippo pathway (Harvey and Hariharan, 2012), as well as *Rfx4, Pdlim5* and *Sox13* (Fig. S4E-F). GO term analysis of the target genes of this network reveled terms related to Wnt and Igf signaling, which control mDA neuron development and promote survival, respectively (Arenas, 2014; Chung et al., 2005; Clarkson et al., 2001), as well as terms related to ECM regulation (Fig. S3G). These results suggest that Rgl3 could influence mDA development by regulating cell signaling and ECM composition in the mDA niche.

### Signaling pathways in the ventral midbrain

To comprehensively characterize signaling patterns within the mDA niche, we compiled ligand-receptor interactions from CellChat (Jin et al., 2021), CellPhoneDB (Efremova et al., 2020), and CellTalkDB (Shao et al., 2021). We then used CellChat, our reference dataset, and scRNA-seq data from the developing VM (La Manno et al., 2016) to predict the primary signaling events in the developing VM. Based on the number of incoming and outgoing interactions, Rgl2,3, pericytes and microglia emerged as major contributors to signaling in the developing VM, with Rgl3 having the most combined interactions (Fig. 4A). Furthermore, analysis of the top predicted signaling interactions revealed pathways known to contribute to mDA neuron development, including both canonical and non-canonical Wnt, Shh, Notch and Fgf (Fig. S6). We also predicted communication related to axon guidance and synapse formation, such as Ntn1-Dcc, Slit-Robo, Ephrin-Eph (Egea and Klein, 2007; Garritsen et al., 2023) and Neurexin-Dag (Gomez et al., 2021), as well as growth factor signals implicated in mDA neuron survival, including the heparin-binding growth factors Pleiotrophin (*Ptn*) (Hida et al., 2007; Moses et al., 2008) and Midkine (*Mdk*) (Ohgake et al., 2009; Prediger et al., 2011) (Fig. S6). Other molecules known to affect mDA differentiation or survival, either directly or indirectly via glial cells, such as the ECM glycoprotein Tenascin-c (*Tnc*), (Garcion et al., 2001; Garwood et al., 2004; Karus et al., 2011; Sousa et al., 2007), and the growth factor *Vegf,* (Falk et al., 2011; Piltonen et al., 2011; Yasuhara et al., 2005) were also identified.

**Figure 4.**
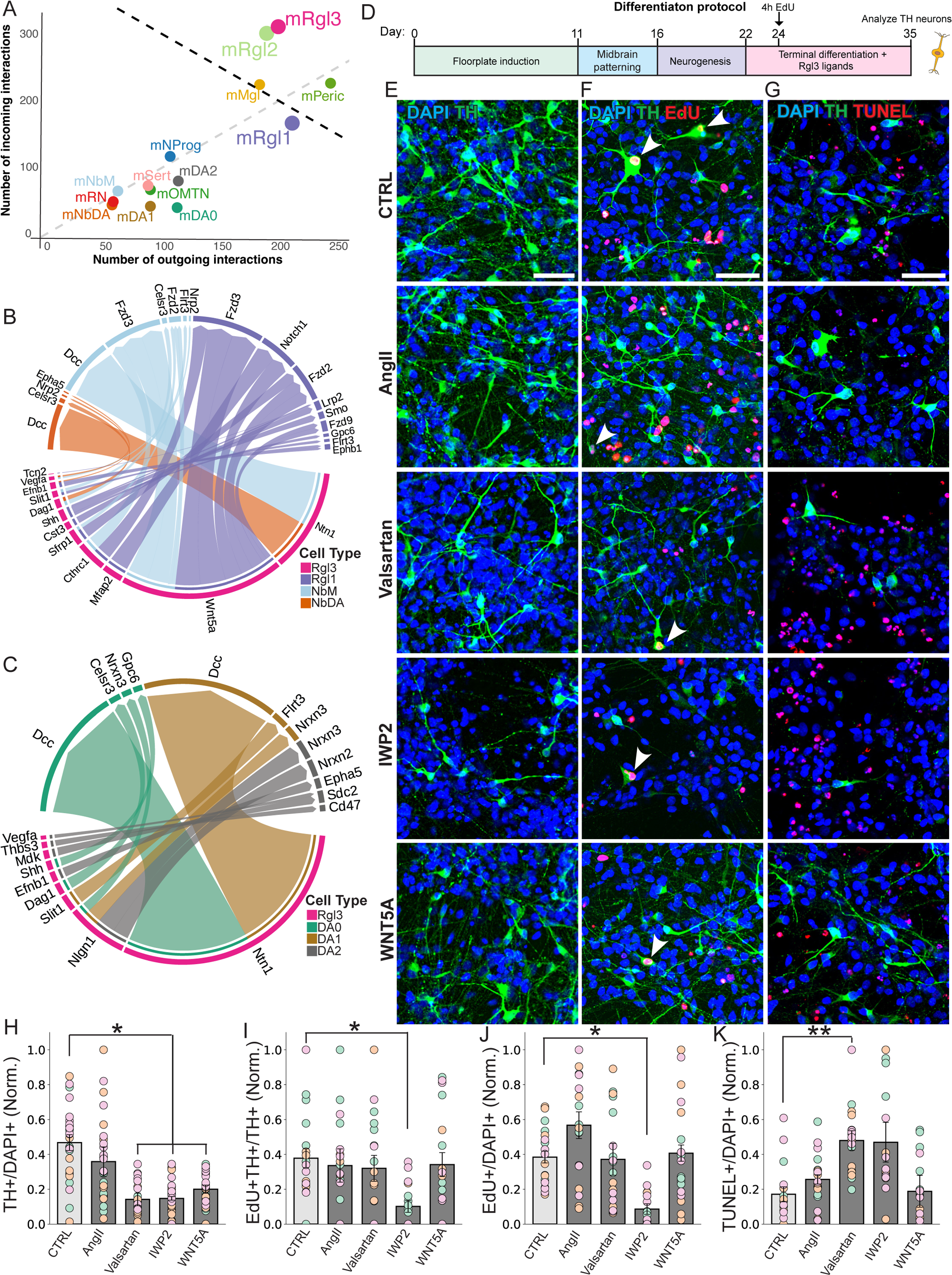
Rgl3 ligands regulate diverse aspects of mDA neuron development. **(A)** Scatter plot showing the number of predicted incoming and outgoing signaling interactions for different VM cell types as predicted by CellChat. Black dotted line represents the 95^th^ percentile of the bootstrapped distribution of the combined incoming and outgoing interaction scores. Grey dotted line (x = y) delineates cell types with more outgoing (above) or incoming (below) signaling interactions. Dots corresponding to Rgl cell types were enlarged for visualization purposes. **(B-C)** Chord diagram showing interactions of Rgl3 with cell types of the dopaminergic lineage as predicted by CellChat. Interactions involve >10 cells. Bar size reflects the probability. Outgoing signals (ligands) from Rgl3 are predicted to interact with receptors expressed by Rgl1, NbM and NbDA (**B**), as well as DA0-2 neurons (**C**). **(D)** Schematic overview of the differentiation protocol used to generate mDA neurons from hESCs. Cells are differentiated in the presence of selected Rgl3 ligands between days 22-35, pulsed with EdU for 4 hours on day 24 to mark proliferating cells, and analyzed at day 35. **(E-G)** Immunostaining showing mDA neurons (TH^+^) (**E**), mDA neurogenesis (TH^+^EdU^+^, arrows in F), proliferation (EdU^+^TH^−^) (**F**), and cell death (TUNEL^+^ nuclei) **(G)** at day 35, after treatment with Rgl3 ligands and signaling molecules. Scale bar in E-G, 100 µm. (**H**) Quantification of TH^+^/DAPI^+^. CTRL: 0.47 ± 0.045; AngII: 0.358 ± 0.084; Valsartan: 0.102 ± 0.036, p = 0.044; IWP2: 0.128 ± 0.010, p = 0.023; WNT5A: 0.174 ± 0.023, p = 0.049. (**I**) Quantification of TH^+^EdU^+^/TH^+^. CTRL: 0.385 ± 0.033; AngII: 0.336 ± 0.096; Valsartan: 0.320 ± 0.074; IWP2: 0.101 ± 0.034, p = 0.048; WNT5A: 0.342 ± 0.068. (I) Quantification of EdU^+^/DAPI^+^. CTRL: 0.385 ± 0.033; AngII: 0.567 ± 0.076; Valsartan: 0.372 ± 0.091; IWP2: 0.086 ± 0.030; WNT5A: 0.407 ± 0.046. (**K**) Quantification of TUNEL^+^/DAPI^+^. CTRL: 0.171 ± 0.057; AngII: 0.256 ± 0.074, Valsartan: 0.480 ± 0.062; IWP2: 0.470 ± 0.122; WNT5A: 0.188 ± 0.094. Quantifications in (H-K) were performed in cells on day 35. Data is normalized to the average of each experiment and values are presented as mean ± SEM with p-values when significant differences were observed. *, p < 0.05, **p < 0.01. Multiple testing correction was performed using Bonferroni-Holm correction. Each data point is represented by a circle, and the circle’s color corresponds to the biological replicate (n = 3 independent experiments).

### Interactions of Rgl3 ligands with cells of the mDA lineage

CellChat analysis of early mDA lineage cell types (Rgl1, NbM, NbDA) revealed that transcripts encoding ligands of the Wnt, Shh, and Notch signaling pathways dominated predicted interactions of Rgl3 with their corresponding receptors in Rgl1. Wnt signaling was also predicted in interactions of Rgl3 with the two neuroblast cell types (NbM, NbDA), but they were mainly characterized by signaling events related to axon guidance, such as Ntn1-Dcc, Efn-Eph, and Dag1-Celsr3 (Lindenmaier et al., 2019) (Fig. 4B). Similarly, Rgl3 was also predicted to interact with the three types of embryonic mDA neurons (DA0-2) through axon guidance molecules, such as Ntn1, and to a lesser degree Slit1, and through synaptic cell adhesion proteins such as Neurexins-Neuroligins (Fig. 4C). These results suggest that Rgl3 ligands may interact with cell types along the mDA lineage and control their development in a cell-type and pathway specific manner.

These predictions prompted us to investigate and experimentally validate whether ligands expressed by Rgl3 could influence human mDA neuron development *in vitro*. We curated a short list of ligands (Fig. S7A) and then selected a few molecules from the Wnt, Bmp, and angiotensin II (AngII) pathways to investigate in human embryonic stem cells (hESCs) (Table 2). A human development-based protocol (Nishimura et al., 2023a) was used to differentiate WA09 cells into cells of the mDA lineage that are comparable to their endogenous counterparts, as assessed by scRNA-seq (Nishimura et al., 2023b). Ligands or small molecules were added to the cultures between days 22 and 35 of differentiation (Fig. 4D), when Rgl3 emerge and mDA progenitors undergo neurogenesis and differentiate into mDA neurons. We then quantified the proportion of TH^+^ neurons generated in the presence of the selected Rgl3 ligands and small molecules (Fig. 4E, 4H, and. S7B-C). While NBL1, BMP1, FGF7, and the AngII receptor type 2 (AGTR2) activator CPG-42112A had no effect, we observed a significant reduction in the proportion of TH^+^ neurons after inhibition of Wnt secretion with IWP2 (73%), or after non-canonical WNT activation with WNT5A (63%) compared to control. However, the most significant decrease (78%) was induced by the AGTR1 inhibitor Valsartan. AGTR1 is particularly interesting because it is expressed in a subtype of human mDA neurons that has recently been associated with PD (Kamath et al., 2022), but the function of AGTR1 in human mDA neuron development had not been previously examined.

**Table 2.**

| Condition | Concentration |
| --- | --- |
| AngII | 100 nM |
| BMP1 | 100 ng/ul |
| NBL1 | 100 ng/ul |
| FGF7 | 100 ng/ul |
| DKK3 | 100 ng/ul |
| WNT5A | 100 ng/ul |
| Valsartan | 10 μM |
| IWP2 | 10 μM |
| CGP-42112A | 300 nM |

### Rgl3-derived ligands regulate different aspects of human mDA neuron development

To investigate the potential cellular mechanisms underlying the decrease in mDA neurons caused by the Wnt and angiotensin pathways, we assessed changes in progenitor proliferation and mDA neurogenesis after an EdU pulse (4h at day 24) to label proliferating cells (Fig. 4D). Notably, treatment with IWP2, unlike WNT5A, ANGII or Valsartan, significantly reduced the proportion of cells undergoing mDA neurogenesis (74%) and proliferation in the culture (78%) (Fig 4F, I, J). These results underline the importance of WNT signaling for human mDA progenitor proliferation and mDA neurogenesis, and of WNT5A for subsequent mDA differentiation as previously described in mice (Andersson et al., 2013).

We also examined the proportion of cells undergoing apoptosis by TUNEL staining and found that Valsartan, but not ANGII, IWP2, or WNT5A, significantly increased (181%) the proportion of TUNEL positive nuclei in the culture (Fig. 4G, K), which together with the lower number of TH+ neurons (Fig. 4H) suggests a new role for the AGTR1 receptor in maintaining the survival of developing mDA neurons. Thus, our analysis demonstrates that different cell types in the mDA lineage are specifically regulated by signaling pathways that can be activated by Rgl3 ligands.

### The extracellular matrix in the ventral midbrain

Our GO analysis of the mDA niche suggested a more significant role for the ECM in mDA neuron development than previously expected. To rank the contribution of each VM cell type to the ECM, we used Matrisome gene sets for core ECM components and ECM regulators (Hynes and Naba, 2012; Naba et al., 2012), and developed a score based on the number of genes expressed and their expression level (see methods). Six cell types (Rgl2,3, ependymal cells, endothelial, pericytes, and microglia) were above a threshold set at the 99.9% quantile of the mean (Fig. 5A) and contributed 82.8% of the mRNA molecules for ECM core components (Fig. S8A), and 88.2% for ECM regulators in the VM (Fig. S8B). The main contributors of ECM core components were pericytes, which express multiple collagen and laminin genes (Fig. S8C) important for ECM structure and organization (Aumailley, 2013; Ricard-Blum, 2011). In contrast, microglia were identified as the main regulators of the ECM, particularly due to their expression of multiple cathepsin proteases (Fig. S8D), which play a role in the immune response (Chapman et al., 1997). Notably, our results identified Rgl3 as the Rgl that contributes the most to the ECM, around 13% of both ECM regulators and ECM core components to the mDA niche (Fig. S8A-B). This included genes commonly expressed by Rgl3 and other VM cell types, such as *Sparc* (also in endothelial cells and pericytes) and *Spon1* (in all Rgl), but also Rgl3-specific regulators, such as *Sulf1* (Fig. 5B) and Rgl3-specific core components, including *Netrin1 (Ntn1)*, *Slit1*, and *Decorin (Dcn)* (Fig 5C). This finding was interesting as our CellChat analysis had predicted interactions of Rgl3 with cells of the mDA lineage involving Ntn1 and Slit1 (Fig. 4B,C), and both factors have previously been implicated in mDA axonal development and guidance (Kastenhuber et al., 2009; Xu et al., 2010). Conversely, *Dcn* has previously been found to regulate midbrain neurogenesis in chicken embryos (Long et al., 2016). This led us to examine the hypothesis that Rgl3 may contribute specialized ECM core components that influence mDA neuron development.

**Figure 5.**
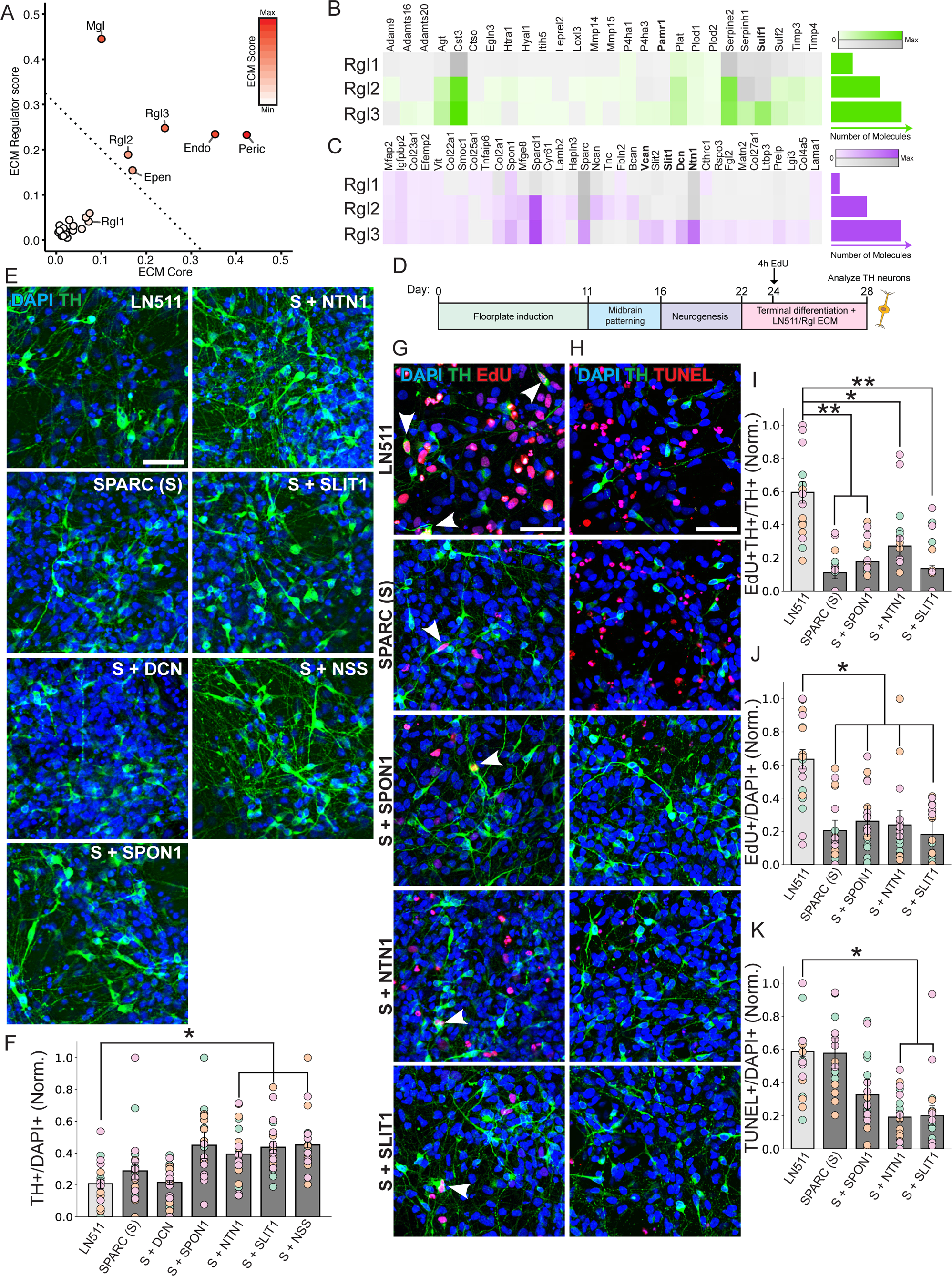
Rgl3 ECM proteins promote survival and increase the number of DA neurons. **(A)** Scatter plot showing the contribution of different VM cell types to the ECM as determined by scores for ECM core components and ECM regulators. Dotted line represents quantile 99.9% of the bootstrapped mean of the ECM scores. **(B, C)** Heat maps showing the expression levels of ECM regulators **(B)** and ECM core components **(C)** in radial glia cell types. Color intensity is proportional to the Bayesian estimate of expression level. Gray scale indicates values below significance level. Bar plot to the right represents total average of transcripts for ECM core components/regulators per cell type. Rgl3 specific ECM regulators and core components are shown in bold. **(D)** Schematic overview of the differentiation protocol used to generate mDA neurons from hESCs. Cells were differentiated in plates coated with Rgl3 ECM core components between days 22-28, pulsed with EdU on day 24 for 4 hours to mark proliferating cells, and analyzed on day 28. **(E)** Immunostaining of TH^+^ neurons on day 28, after treatment with ECM proteins. NSS = NTN1 + SLIT1 + SPON1. Laminin-511 (LN511) was used as control. Scale bar, 100 µm. **(F)** Quantification of TH^+^/DAPI^+^. LN511: 0.207 ± 0.029; SPARC: 0.288 ± 0.052; S + NTN1: 0.393 ± 0.018, p = 0.035; S + SLIT1: 0.437 ± 0;039, p = 0.047; S + SPON1: 0.449 ± 0.084; S + NSS: 0.452 ± 0.019, p = 0.015. Data is normalized to the average of each experiment and values are presented as mean ± SEM with p-values when significant differences were observed. *, p < 0.05. Multiple testing correction was performed using Bonferroni-Holm correction. Each data point is represented by a circle, and the circle’s color corresponds to the biological replicate (n = 3 independent experiments). **(G-H)** mDA neurogenesis (TH^+^EdU^+^, arrowheads) **(G)**, Proliferation (EdU^+^TH^−^) **(G)** and cell death (TUNEL^+^ nuclei) **(H)** were examined by immunocytochemistry on day 28 after cultivation on selected Rgl3 ECM proteins. A 4-hour EdU pulse was performed on day 24. Scale bar, 100 µm. **(I)** Quantification of TH^+^EdU^+^/TH^+^. LN511: 0.594 ± 0.064; SPARC: 0.110 ± 0.036, p = 0.009; S + NTN1: 0.270 ± 0.061, p = 0.020; S + SLIT1: 0.135 ± 0.012, p = 0.0095; S + SPON1: 0.179 ± 0.003, p = 0.0095. **(J)**, EdU^+^/DAPI^+^. LN511: 0.635 ± 0.058; SPARC: 0.205 ± 0.062, p = 0.029; S + NTN1: 0.238 ± 0.088, p = 0.044; S + SLIT1: 0.181 ± 0.090, p = 0.041; S + SPON1: 0.261 ± 0.094, p = 0.044. **(K)** TUNEL^+^/DAPI^+^. LN511: 0.585 ± 0.029; SPARC: 0.576 ± 0.037; S + NTN1: 0.185 ± 0.024, p = 0.002; S + SLIT1: 0.2 ± 0.060, p = 0.014; S + SPON1: 0.327 ± 0.173. Quantifications in (I-K) performed on day 28. Data is normalized to the average of each experiment and values are presented as mean ± SEM with p-values when significant differences were observed. *, p < 0.05, **p < 0.01. Multiple testing correction was performed using Bonferroni-Holm correction. Each data point is represented by a circle, and the circle’s color corresponds to the biological replicate (n = 3 independent experiments).

### Rgl3-specific ECM proteins interact with and regulate the development of human midbrain cells of the DA lineage

To investigate the function of Rgl3 ECM proteins in mDA neuron development, we selected three ECM proteins unique to Rgl3 (Ntn1, Slit1, and Dcn), one common to all Rgl (Spon1) and one found in Rgl3, pericytes and endothelial cells (Sparc). hESCs were differentiated towards mDA neurons in plates coated with laminin 511 (LN511, as control), which is known to promote mDA neuron differentiation and survival (Zhang et al., 2017), or the Rgl ECM proteins, from day 22-28 (Fig. 5D). While SPARC *per se* did not change the proportion of TH^+^ neurons compared to LN511 (Fig. 5E,F), the combination of SPARC with the two Rgl3-specific ECM proteins predicted by CellChat, NTN1 or SLIT1, increased the proportion of TH^+^ neurons in the culture by 90% and 111%, respectively (Fig. 5E-F). However, their effects together with SPON1 (NSS) were not additive, suggesting that they may work through a similar mechanism. Notably, neither SPON1 (common Rgl) nor DCN (Rgl3-specific but not predicted by CellChat), significantly increased the yield of mDA neurons (Figure 5E,F), indicating that of the proteins tested, only two Rgl3-specific ECM proteins predicted by CellChat, NTN1 and SLIT1, control the development of mDA neurons *in vitro*.

To investigate the cellular mechanism by which the ECM proteins increase the yield of DA neurons, we examined progenitor proliferation (EdU^+^/DAPI^+^) and mDA neurogenesis (EdU^+^TH^+^/TH^+^) from day 24-28. Compared to LN511, SPARC decreased proliferation and neurogenesis in all conditions (Fig. 5G,I-J), indicating an anti-proliferative and anti-neurogenic effect on midbrain non-DA cells, as reported for other cell types (Francki et al., 2003; Schiemann et al., 2003). We next determined whether NTN1 or SLIT1 could increase the number of mDA neurons by preventing cell loss and found they reduced the proportion of TUNEL^+^ cells compared to control by 68% and 66%, respectively (day 22-28, Fig. 5D,H,K), while no changes were induced by SPARC or SPON1. Thus, our results suggest a novel function of NTN1 and SLIT1 in preventing cell death and enhancing the yield of TH^+^ neurons.

### Roles of Rgl1 and Rgl3 in endogenous human ventral midbrain development

To investigate whether Rgl1 and Rgl3 have similar specialized roles in human midbrain development *in vivo*, we used a recent scRNA-seq atlas of the developing human brain, which covers the development of the VM during the period of mDA neurogenesis (5-14 weeks post conception) (Braun et al., 2023). All three Rgl cell-types (hRgl1-3) were detected in the human data set, and GSEA analysis confirmed their similarity to mouse Rgl1-3 (mRgl1-3) (Fig. S9A-C). VM cells segregated into three major axes of differentiation, with one axis characterized by basal plate markers (*NKX6-1*, *NKX6-2*, or *NKX2-2*^+^) and another by FP markers (*FOXA2^+^/LMX1A^+^/EN1^+^*) (Fig. 6A,B). Based on this classification, Rgl1 were identified in the FP, Rgl2 in the basal plate, and Rgl3 in both regions. Moreover, Rgl1 cells were organized into two main clusters, one in the posterior FP (*FOXA2*^+^*/LMX1A*^+^*/EN1*^+^), which gives rise to mDA neurons (*, Fig. 6A, B), and a second cluster expressing the anterior marker *PITX2* (Braun et al., 2023), found in subthalamic nucleus neurons (Kee et al., 2017).

**Figure 6.**
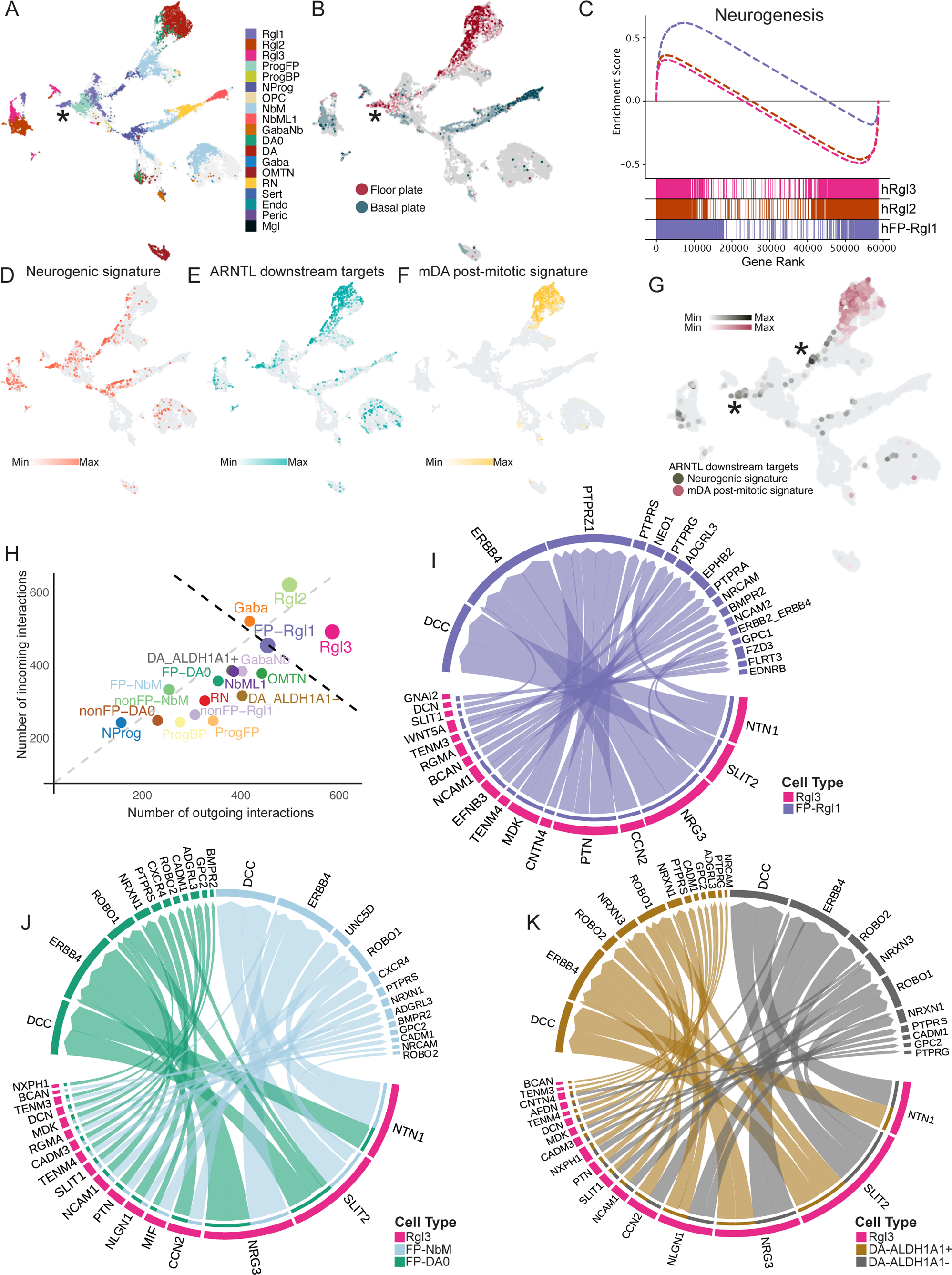
Analysis of endogenous human Rgl1 and Rgl3 *in vivo,* in the developing ventral midbrain. **(A)** UMAP of cells in the developing human ventral midbrain (*FOXA2*/*FOXA1*^+^ or *TH*^+^) between weeks 5-14 post conception, as described in Braun et al. 2023. Cell type identity was predicted by logistic regression trained on human VM cell types as defined in La Manno et al. 2016. Colored cells, probability >60%; gray cells, probability <60%. **(B)** UMAP of cells colored based on enriched floor plate (*FOXA2*^+^/*LMX1A*^+^/*EN1*^+^) or basal plate (*NKX2.2^+^*, *NKX6.1^+^* or *NKX6.2^+^)* annotation. **(C)** Gene set enrichment analysis (GSEA) of human Rgl1, 2 and, 3 transcriptomes using human neurogenesis gene set (GO: 0022008). Floor plate Rgl1 was isolated from non-floor plate Rgl1 by the expression of genes defining floor plate identity (*LMX1A+FOXA2+EN1+)*. FP-Rgl1: NES = 1.074, FDR = 0.068. Rgl2: NES = −1.072, FDR = 0.198. Rgl3: NES = −1.107, FDR = 0.116. **(D)** UMAP of VM cells showing the expression of the five most significant genes in the neurogenic signature defined in mouse Rgl1 (*ARNTL, E2F3, E2F5, ASCL1, SOX5* as defined in 2C) in addition to *NEUROG2.* Color intensity corresponds to expression level. Cells are colored if >3 genes of the neurogenic signature are expressed in the same cell. **(E)** UMAP of VM cells showing the expression of genes in the ARNTL downstream target gene signature (*BMP7*, *DVL2*, *NEUROD1*, *GSK3B*, *TCF12*, *NR4A2*, *HDAC2*, *IGF1R*, as defined in Fig. 2G). Color intensity corresponds to expression level. Cells are colored if >5 genes of the ARNTL downstream signature are expressed in the same cell. **(F)** UMAP of VM cells showing the expression of a mDA post-mitotic signature (*EN1*, *NR4A2*, *PITX3*, *TH*). Color intensity corresponds to expression level. Cells are colored if all four genes are expressed in the same cell. **(G)** UMAP of VM cells showing shared expression of the ARNTL downstream target gene signature (E) and the neurogenic (D) or the mDA post-mitotic signature (F). Color intensity corresponds to expression level. **(H)** Scatter plot showing the number of predicted incoming and outgoing signaling interactions for the different VM cell types found in the human developing VM, as predicted by CellChat. Cells of the DA lineage expressing markers of posterior midbrain identity (*LMX1A^+^/FOXA2^+^/EN1^+^*) were defined as FP. Black dotted line represents the 95th percentile of the bootstrapped distribution of the combined incoming and outgoing interaction scores and the gray dotted line (x = y) delineates cell types with more outgoing (above) or incoming (below) signaling interactions. Dots corresponding to Rgl cell types were enlarged for visualization purposes. **(I-K)** Chord diagram showing predicted interactions of Rgl3 with floor plate: Rgl1 (**I**), NbM and DA0 (**J**), and DA-ALDH1A1^+^ and DA-ALDH1A1^−^ (**K**). Interactions were filtered to a minimum communication probability of 3%.

GSEA analysis of hRgl1-3 identified hFP-Rgl1 (*FOXA2^+^/LMX1A^+^/EN1^+^*) as the primary contributor to neurogenesis and indicated that genes in the mRgl1 transcriptional network, such as *ASCL1*, *TCF3/4/12*, and *SOX5*, also contribute to neurogenesis in hFP-Rgl1 (Fig. 6C and Table S3). Moreover, GSEA identified novel neurogenesis-related genes in hFP-Rgl1 previously shown to enrich for mDA progenitors, including *TPBG* and *PTPRO* (Xu et al., 2022; Yoo et al., 2021).

To explore a potential role of the *ARNTL*-centered mRgl1 neurogenic transcriptional program in hFP-Rgl1, we utilized the *ARNTL* downstream target gene signature (Figure 2G), and defined a neurogenic signature composed of TFs from the mRgl1 neurogenic network (Fig. 2C,E,F) and *NEUROG2*, a TF required for mDA neurogenesis (Kele et al., 2006). Analysis of the human mDA lineage revealed two cell types exhibiting both signatures: hFP-Rgl1 and late-stage NProg (*, Fig. 6D and E), indicating that neurogenesis is activated along the Rgl1-NProg axis of differentiation. Moreover, the *ARNTL* downstream signature was expressed in embryonic mDA neurons (*EN1^+^/NR4A2^+^/PITX3^+^/TH^+^*) (Fig. 6F). Combined, our results suggest that a neurogenic program contributed by ARNTL is initiated in hFP-Rgl1, maintained in NProg, and ends in mDA neurons (Fig. 6G).

### Human Rgl3 as a signaling center in the human developing ventral midbrain

CellChat analysis identified Rgl3 as the cell type with the most outgoing signaling interactions in the human developing VM (Fig. 6H), suggesting that Rgl3 may also act as a signaling center in human VM development. Rgl3 was predicted to interact with all cells of the DA lineage in the posterior mFP (*LMX1A^+^/FOXA2^+^/EN1^+^*) (Fig. 6B). Surprisingly, these interactions were more diverse but less distinct between cell types, compared to mice (Fig. 4). Throughout the mDA lineage, the majority of predicted Rgl3 interactions involved ligands important for axon guidance, cell migration, and cell survival, such as *NTN1*, *SLIT1/2, NRG3, NCAM1, PTN, MDK,* etc. (Fig 6I-K) (Bartolini et al., 2017; Garritsen et al., 2023; Klein and Pasterkamp, 2021; Lin et al., 2005; Ohgake et al., 2009, 2009; Rahman-Enyart et al., 2020; Schmid and Maness, 2008; Xu et al., 2010). The most common interactions were mediated by *NTN1* and *SLIT2* (Fig 6I-K), but they involved distinct receptors in pre- and post-mitotic cells, suggesting possible cell type specific signal transduction. While *NTN1* was predicted to interact with *NEO* in FP-Rgl1 and *UNC5D* in FP-NbM and FP-DA0; *SLIT1/2* involved *GPC1/FLRT3* in FP-Rgl1 and *ROBO* in FP-NbM and FP-DA0. Other hRgl3 factors predicted to interact with FP-Rgl1 included *WNT5A* (Fig. 6I), but not *SHH* or *NOTCH*, two factors detected in mRgl3 (Fig. 4B). These findings may be due to species differences but may also reflect the time at which Rgl1 was captured (week 7-8 in human vs E11.5-12.5 in mice Rgl1, ≈ week 6-7 in human, (Clancy et al., 2001) (Fig. S9D-G).

Additionally, Rgl3 was predicted to interact with the postmitotic NbM through *MIF* and the chemokine receptor *CXCR4*, which regulates the migration of mDA neuroblasts (Bodea et al., 2014; Yang et al., 2013), and through *NLGNs* with *NRXNs* and *CADM*, which are involved in synapse formation (Biederer et al., 2002) (Fig. 6J). Finally, Rgl3 was predicted to interact with two embryonic DA neuron subtypes, defined by *ALDH1A1* expression (Fig. S9H). Selective interactions predicted for DA-ALDH1A1^−^ and DA-ALDH1A1^+^ neurons involved *TENM3/4-ADGRL3* and *NRXN1-LRRTM4*, respectively (Fig. 6K), suggesting distinct regulation of synapse organization (Roppongi et al., 2020; Zhang et al., 2022). Additionally, TENM4 is involved in axon guidance, and loss-of-function and missense variants of this gene are associated with essential tremor and early onset PD (Hor et al., 2015; Pu et al., 2020).

Thus, our *in silico* analysis of the human VM *in vivo* validates our *in vitro* functional analysis demonstrating that Rgl3 factors regulate multiple aspects of human mDA neuron development, and suggests a function of Rgl3 as a signaling center in the mDA niche.

## Discussion

Rgl cells are a group of heterogeneous CNS cell types, with divergent and poorly understood roles at the individual level. Here we report that Rgl cell types serve highly specialized and distinct functions that are largely conserved from mouse to human. Our single-cell transcriptomics and functional analysis define mouse and human Rgl1 as mDA progenitors and Rgl3 as the main midbrain-specific signaling component of the mDA niche. Moreover, our study defines a neurogenic network in Rgl1 that controls mDA neurogenesis and is centered on the TF *ARNTL*, a novel cell intrinsic regulator of mDA neuron development. Additionally, we uncover multiple cell extrinsic factors expressed by Rgl3, including morphogens (WNT5A), signaling factors (ANGII) and ECM molecules (SPARC, NTN1 and SLIT1), which control the proliferation and neurogenesis of mDA progenitors as well as the differentiation and survival of human mDA neurons.

Rgl1 was identified as a neurogenic cell type in both species. TFs commonly enriched included *Ascl1* (Kele et al., 2006) and *Tcfs* (Mesman and Smidt, 2017), which regulate mDA neurogenesis, as well as *Sox5*, which regulates neural cell fate decisions (Lai et al., 2008). The mRgl1 transcriptional profile was regulated by a pro-neurogenic TF network centered on *Arntl/Bmal1,* a TF which we demonstrate can regulate the timing of cell cycle exit and mDA neurogenesis in hLT-NES cells (Fig. 3). Moreover, we find that hRgl1 is enriched in *NEUROG2* and TFs of the mRgl1 *ARNTL* neurogenic network and *ARNTL* target genes are enriched in endogenous human mDA neurons. Since *ARNTL* promotes mDA neurogenesis *in vitro*, our findings suggest that *ARNTL* regulates mDA neurogenesis in midbrain development. Combined, our results define Rgl1 as a neurogenic mFP Rgl subtype and as a mDA progenitor.

Conversely, our analysis of Rgl3 suggested a role for this cell type as a signaling center in the developing VM. mRgl3 was defined by a transcriptional network centered on *Tead1*, which induces *Shh* and *Foxa2* expression in the FP through interactions with *Yap* (Cheng et al., 2023; Sawada et al., 2008). Moreover, the second most significant TF in this network was *Rfx4*, which also regulates *Shh* expression and controls ventral midline formation (Ashique et al., 2009; Sedykh et al., 2018). Notably, mutation of *Tead1* or *Rfx4* cause dorsoventral patterning defects (Cheng et al., 2023; Sedykh et al., 2018), suggesting an important function of Rgl3 TFs in delineating the FP midline and specifying ventral identities. Accordingly, CellChat defined Rgl3 as the cell type with most outgoing predicted signaling interactions, involving all cell types of the mDA lineage. Common factors predicted in mouse and human Rgl3 included morphogens and modulators of the Wnt pathway, axon guiding molecules (Netrin1, Slits) and multiple ECM proteins, all of which are traditional mFP-derived factors (Arenas et al., 2015; Garritsen et al., 2023).

Analysis of the function of Rgl3 signals at the time when Rgl3 emerge from hPSCs during mDA differentiation (Nishimura et al., 2023b) revealed new functions in the mDA lineage. We found that inhibition of Wnt secretion with IWP2 reduced progenitor proliferation, mDA neurogenesis, and the number of mDA neurons, uncovering the necessity of Wnt signaling for maintenance of human mDA neuron development. Additionally, we found that inhibition of the angiotensin receptor AGTR1 caused cell loss and decreased the number of TH^+^ cells, suggesting a novel function of endogenous Rgl3-derived AngII in maintaining the survival of human embryonic mDA neurons. This finding is particularly interesting since *AGTR1+* adult mDA neurons in the human substantia nigra are more vulnerable to PD (Kamath et al., 2022), and the angiotensin pathway has been implicated in PD pathology (Mertens et al., 2010; Zawada et al., 2015), and in modulating DA neurotransmission (Brown et al., 1996; Medelsohn et al., 1993; Villar-Cheda et al., 2010). Thus, our results uncover a novel activity of AngII in promoting human mDA neuron survival, which might be of interest in the context of neurodegeneration. Moreover, we demonstrate that two ECM proteins uniquely expressed by Rgl3, NTN1 and SLIT1, prevent cell death and increase the number of mDA neurons. While Ntn1 is known to promote mDA neuron survival in the adult mouse substantia nigra (Ahn et al., 2020; Jasmin et al., 2021; Lo et al., 2022), no survival-promoting effects had been described for NTN1 or SLIT1 in human embryonic development. Thus, our analysis uncovers novel activities of distinct Rgl3 extrinsic factors, such as ligands and ECM proteins, which control multiple aspects of human mDA neuron development, as predicted *in vitro* and *in vivo*. These results point to Rgl3 as a critical component of the signaling center in the developing mFP.

Taken together, our *in vitro* and *in vivo* results shed light on the cell types and factors contributing to the mDA niche and uncover highly specialized functions of mouse and human Rgl subtypes within the mFP. Our data suggest a model in which Rgl1, as a neuronal progenitor, and Rgl3, as a signaling cell, form a developmental unit capable of generating and sculpturing the phenotype of mDA neurons. Moreover, we provide evidence that both cell-intrinsic Rgl1 and cell-extrinsic Rgl3-derived factors can improve the generation of human mDA neurons from stem cells, suggesting a possible application of these factors to address current challenges (Kim et al., 2020) and to improve regenerative medicine approaches for the treatment of PD.

## Materials and methods

### Bulk transcriptome determination

Mouse embryos were obtained from TH-GFP animals (Matsushita et al. 2002) that were mated overnight, and noon of the day the plug was considered E0.5. Embryos were dissected out of the uterine horns at E11.5-E14.5 and placed in ice-cold sterile PBS where brain regions were dissected under a stereomicroscope with a UV attachment to detect GFP. Tissue samples were collected in separate tubes, stored at −80°C until RNA isolation. Ethical approval for mice experimentation was granted by the local ethics committee, Stockholm Norra Djurförsöksetiska Nämnd number N326/12 and N158/15.

Total RNA isolation was performed with RNeasy kit (Qiagen). The RNA integrity and concentration was checked using Qubit and 2200 TapeStation (Agilent). Illumina TruSeq libraries were prepared using kit and protocols from Illumina. High-throughput sequencing was performed on a HiSeq 2000.

### Bulk Genes Expression Analysis

Differentially expressed genes (DEG) on each developmental stage were identified utilizing Qlucore Omics Explorer v3.1 (Qlucore AB, Lund, Sweden) utilizing t-test comparing VM samples versus other regions, with false discovery rate q-value correction (Storey, 2002). Variance filter of 15% and a threshold for significance of p-value = 0.01 across stages, unless otherwise stated. Sample correlation was calculated from log2(RPKM+1) values after variance filtering of 12.5%. Significant expression is baseline (>99.8% posterior probability). Analyses were made using R statistical software version 3.4.0 and ggplot2 version 2.2.1 (Wickham, 2016) for plots. RNA-seq data are deposited in the public NCBI GEO database with accession number GSE82099 for E12.5 (Gyllborg et al., 2018) and GSE117394 for E11.5, E13.5 and E14.5.

### Bulk Gene Expression Analysis Weighted Gene Co-Expression Network Analysis

(WGCNA) (Langfelder and Horvath, 2008) was performed with the log2(RPKM+1) transformed values, filtered by variance until 10,068 genes were selected. The topological overlap matrix was calculated with the variables of a signed network, with power of 7, as this is the lowest power that results in a scale free network (Horvath, 2011). The identification of modules was performed with the “tree” option on a minimum module size of 100; modules with correlation higher than 90% were merged. Module enrichment for DEG in VM at all analyzed stages was done with Fisher’s exact test with false discovery rate q-value correction (Storey 2002). A score for the enrichment was calculated as the product between the –log10(q-value) and a standardized z-score for DEG per module. Module network layout was made using Cytoscape and interactions in the top 5% of correlation (from WGCNA) were selected for further analysis. Expression changes over time are represented by the color of each node, which was calculated as the normalized difference of RPKM between late (E13.5, E14.5) and early (E11.5, E12.5) for each gene.

The identification of VM gene modules with developmental or stage dependent expression were identified with WGCNA on the VM samples from E11.5 to E14.5. Further filtering for genes not expressed in the VM was performed until 9,061 genes were selected. The topological overlap matrix was calculated with the options of a signed network, with power of 17. The identification of modules was performed with the “tree” option on a minimum module size of 100 genes, modules with correlation higher than 99% were merged. Correlation with samples trait and Student asymptotic p-values were calculated as described (Langfelder and Horvath 2008). Embryonic stage (E11.5 to E14.5) was used as sample trait, ordinal values for stage (1 for E11.5, 2 for E12.5 and so on), and binary values for middle stages (E12.5, E13.5 as 1). Network layouts and analysis, were made using Cytoscape v3.3.0 (Shannon et al., 2003) or Gephi v0.9.1 (Bastian et al., 2009). WGCNA were carried out with the R package v1.49 (Langfelder and Horvath 2008).

### Network single-cell deconvolution

Using data from La Manno et al., each gene in the network was assigned to the cell types that expressed that gene, as identified above baseline threshold (see above and La Manno et al. 2016). Cell types represented by one gene, or cell types with a final contribution fewer than 1% of the genes were excluded from the network.

### Single Cell Gene enrichment Analysis (GSEA)

GSEA was done by pre-ranking the genes by the class difference for the cell type of interest. The analysis was performed on the GSEA software v2.2.2 (Subramanian et al. 2005) with the MSigDB genesets v5.0 for canonical pathways and GO biological processes (Subramanian et al, 2005) with 1000 permutations. Due to the nature of the single-cell transcriptome profiles and molecular counting with unique molecular identifiers (Islam et al., 2014) negative enrichment score (ES) on a gene set for a cell type in particular were interpreted as enrichment on another compared cell type. Analysis of Gene Ontology (GO) enrichment were made utilizing the R package ClusterProfiler (Yu et al., 2012) or MSigDB (Subramanian et al, 2005) with false discovery rate q-value correction (Storey 2002). Enrichment of the targets of TF were analyzed with hypergeometric test and with false discovery rate q-value correction (Storey 2002) over the MSigDB C2 gene sets (Subramanian et al, 2005).

GSEA of human Rgl cell types: mRgl1-3 gene sets were defined by selecting all genes expressed above baseline (>99.8% posterior probability) in mRgl1-3 and then selecting only those genes uniquely expressed by each cell type. Genes related to sex, cell cycle, blood, mitochondria, and immediate early genes were removed from human Rgl transcriptomes prior to the analysis. Analysis was performed using GSEApy (Fang et al., 2023) with pre-rank module with 1000 permutations. The pre-rank input was obtained from performing DEG between human Rgl cell types using scanpy’s rank_genes_groups with method = ‘wilcoxon’ on log-normalized expression matrix (Wolf et al., 2018).

### Single-Cell Transcription Factor (TF) pattern mining

TFs with enriched expression in each radial glia cell type were used as input to obtain the target of the human homologous TF from iRegulon plugin for Cytoscape (Janky et al, 2014), retrieving up to 1,000 target genes per TF. Targets of TFs by cell type detailed on table S3. Clustering of the TF was done by Jaccard index, as an indicator of shared target genes.

Analysis of enrichment of combinatorial TF target genes were done using Fast Westfall-Young (FWY) permutation procedure v1.0.1 (Terada et al, 2013a, 2013b), utilizing a significance threshold of 0.05 in the one-tailed Fisher test and tested with 1,000 permutations. Gene expression among the radial glia cells types was scored using the logarithmic difference between the groups (Subramanian et al, 2005). Values above 0.5 (or 50% of upregulation) were considered enriched in that cell type. The resulting significant combinatorial patterns were represented as a network on which the edge between a set of TFs is measured by an interaction score. This interaction score is calculated for each TF pair as the sum of –log10 (adjusted p-value) for all the TF combinations of that pair. Adjusted p-values <0.001 were consider equal to 0.001 for calculations of interaction score. A control null distribution of transcription enrichment was performed with randomly selected TFs from MSigDB C3 database (Subramanian et al, 2005). Selecting the same numbers of TFs used before for Rgl1-3 respectively. On each cell type 100 random TFs selections were analyzed by FWY maintaining all other settings.

### CellChat Analysis

To investigate potential interactions between cell types in the mouse and human ventral midbrain, cell-to-cell communication analysis was performed using CellChat v1 on the datasets published by La Manno et al. 2016 and Braun et al. 2023. To enrich neurodevelopmentally relevant ligand-receptor pairs, the CellChat ligand-receptor database was merged with the databases of CellPhoneDB (Efremova et al., 2020) and CellTalkDB (Shao et al., 2021). This extended database assumes conservation of ligand-receptor pairs between mouse and human and simplifies some multi-subunit and antagonist interactions modeled by CellChat, as these are not available for all pairs in the extended database. For the mouse dataset, we combined all time points in a single analysis, as the number of cells in the dataset is limited, disregarding the spatiotemporal proximity of celltypes. The mouse dataset (excluding unannotated and endothelial cells) was analyzed following the authors’ workflow: (identifyOverExpressedGenes, identifyOverExpressedInteractions, projectData, computeCommunProb, filterCommunication, subsetCommunication, netAnalysis_computeCentrality), with min.ncells = 10.

To avoid calling signaling activity by lowly expressed genes, gene expression was filtered as in La Manno et al. 2016, to retain only those genes expressed above baseline for any given cell type independently in each cell type. The number of outgoing interactions was calculated by counting the ligand-receptor pairs for which a given cell type is a source and the number of incoming interactions by counting the ligand-receptor pairs for which a given cell type is a target. Before plotting chord plots, we manually removed the ligand-receptor pair the following interactions: Wnt5a-Notch1, and all interactions involving App, Adam10, Adam17, Mmp9, Tyrobp, Ntng, as the support in the literature for these interactions is limited. The CellChat plotting functions were also modified to allow for coloring chord plot arrows by target cell type instead of source cell type. To plot the heatmaps, we aggregated ligand-receptor pairs into single families (Notch, Dlk, Fgf, Igf2, Nrxn, Slit, Sema) by summing communication probabilities.

For the human data, we analyzed cells with midbrain regional annotation (see Braun et al. 2023 for details) following the same steps as above, without filtering based on baseline gene expression. Instead, we prioritized top interactions by thresholding with a minimum communication probability of 3%. To isolate cell types with a floor plate annotation, cells were scored positively based on *LMX1A/FOXA2/EN1* expression and negatively based on *PITX2/NKX6-1/NKX2-2* expression, choosing hard thresholds consistent with the clusters in UMAP space. The 95^th^ percentile of the bootstrapped distribution of the combined incoming and outgoing interaction scores was calculated by removing the source and target components for each signaling axis (represented as source | target | ligand | receptor) identified by CellChat and repopulating the source and target by selecting a cell type from the same tercile of ligand/receptor expression, using a random uniform distribution. This process was iteratively performed 10.000 times for all source | target | ligand | receptor rows, generating bootstrap replicas. Cell type occurrences as both source and target were tallied for each bootstrap replica, producing a compound score which reflects the overall centrality in signaling.

The code and datasets can be found on Github: https://github.com/lamanno-epfl/rgl3_signaling_dopaminergic_analysis/tree/main

### ECM cell type score

Cell type contribution to ECM was calculated with scRNA-seq data from La Manno *et al*, 2016. ECM score was calculated based on the expression of ECM core genes and genes contributing to ECM regulation as defined by the Matrisome project (Naba et al., 2012).

The score for a gene set is calculated as:

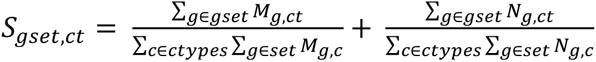

Let *M_g,ct_* be the Bayesian estimate of the expression level of the gene *g* in cell type *ct* and let *N_g,ct_* be the value of the indicator vector that is 1 when the gene *g* is expressed in cell type *ct* above baseline and 0 otherwise. The score is an indicator between diversity of genes in a biological function or gene set and the expression levels of those genes. A combined score by the sum of both components give us an ECM score. The threshold line was set at the percentile 99.9% of the bootstrapped distribution of the mean of the combined score with 10e5 replicates.

### Analysis of human scRNA-seq data

Analysis of human developing ventral midbrain were performed on the dataset published by Braun et al. 2023. For information on how cell type identity was assigned based on the cell types as defined by La Manno et al. (2016), see Braun et al. (2023). To analyze the neurogenic potential of floorplate Rgl1, we only considered the expression of neurogenic genes in Rgl1 with floorplate (*LMX1A+FOXA2+EN1+)* identity. For plotting, the log expression of the product of any combination of 3 or more neurogenic genes (*ARNTL*, *E2F3, E2F5, ASCL1, NEUROG2* and *SOX5*), 5 ARNTL downstream targets (*BMP7, DVL2, NEUROD1, GSK3B, TCF12, NR4A2, HDAC* and*, IGF1R*), and all 4 post-mitotic mDA neuron genes (*EN1+,NR4A2+,TH+* and *PITX3+*) were plotted onto the UMAP.

### Human neuroepithelial stem cell differentiation

Sai2 hLT-NES cells were maintained as previously described for hLT-NES cell (Tailor et al., 2013). For differentiation, hLT-NES cells were dissociated (TrypLE Select, Thermo Fisher Scientific, 12563011) and seeded at a density of 150.000 cells/cm^2^ on glass coverslips coated with poly-L-ornithine (0.02 mg/ml, Sigma, P4957) and laminin (2μg/cm2, Invitrogen, 23017-015). Cells were cultivated in DMEM/F12 (Gibco, 31331028) with 10% FBS. On days 0-1, N2 (1:100, Gibco, 17502001), B27 (1:1,000, Gibco, 17504001), SHH (200 ng/ml, R&D, 461-SH-025), CT99021 (1 μM, Sigma, SML1046) and WNT5A (100 ng/ml, R&D, 645-WN-010) were added to the culture media. The concentration of B27 was increased (1:100) between days 2-3. On days 4-5, culture media was supplemented with N2 (1:100), B27 (1:100), SHH (50 ng/ml), GDNF (20 ng/ml, R&D, 212-GD-050), and WNT5A (100 ng/ml, R&D, 645-WN-010). On days 6-7, B27 and SHH were removed from the culture media and BDNF (20ng/ml, R&D, 248-BDB-050), Ascorbic Acid (200 µM, Sigma, A4403), TGFβ3 (1ng/ml, R&D, 243-B3-010) and dbcAMP (0.5 mM, Sigma, D0627) were added.

### ARNTL loss and gain of function

For ARNTL KD, commercially available ARNTL shRNA lentiviral particles targeted against the human transcript were used (sc-38165, Santa Cruz Biotech.) to generate stable cell lines selected with puromycin (500 ng/ml first 4 days, maintained with 200 ng/ml). As negative control, lentiviral particles against no know human mRNA were used (sc-108080, Santa Cruz Biotech.). Lentiviral production for over-expression, was performed as previously described (Rivetti Di Val Cervo et al., 2017; Villaescusa et al., 2016). ARNTL KD was confirmed with western blot analysis of total cell lysates using anti-Arntl (1:1000, ab93806, Abcam). For loading control, anti-Lamin-B1 was used (1:5000, ab16048, Abcam). Differentiation experiments were performed using stable shRNA cells with less than 10 passages counted from the lentiviral infection.

For ARNTL OE, the coding sequence of mouse ARNTL was synthesized (Twist Biosciences) and cloned into the NheI site of FUW-TetO-MCS (Addgene plasmid #84008, kindly donated from Professor Stefano Piccolo). The final plasmid was verified by sequencing before usage. The Tet-O-FUW-EGFP was obtained from Addgene (#30130). Lentiviral production was performed as previously described (Mattiassi et al., 2021). ARNTL OE was verified by immunocytochemistry with anti-Arntl (1:1000, ab93806, Abcam). ARNTL expression was induced during differentiation using doxycycline (Sigma, 2 µg/ml for day 4 and 1 µg/ml for day 5).

### Human embryonic stem cell differentiation

WA09 hESCs were obtained from WiCell, maintained, and differentiated as previously described (Nishimura et al., 2023b), with the addition of SHH-C24II (200 ng/ml, Miltenyi Biotec, 130-095) between days 7 and 10 of the differentiation protocol. On day 22, cells were prepared for ligand and ECM testing. Cells were dissociated (TrypLE Select) and seeded at a density of 500.000 cells/cm^2^ in 96 well plates. For ligand testing, cells were cultured with WNT5A (100 ng/ml, R&D, 645-WN-010), IWP2 (10 µM, Tocris, 3533), Valsartan (10 µM, Sigma-Aldrich, SML0142), AngII (100 nM, Sigma-Aldrich, A9525), BMP1 (100 ng/ml, R&D, 1927-ZN-010), FGF7 (100ng/ml, R&D, 251-KG), NBL1 (100ng/ml, R&D, 755-DA-050), DKK3 (100ng/ml, R&D, 1118-DK-050) or CGP-42112A (300 nM, Sigma-Aldrich, C160) between days 22 and 35. For ECM experiments, plates were coated with SPARC (R&D, 941-SP-050), SPARCL1 (R&D, 2728-SL-050), NTN1 (R&D, 6419-N1-025), SLIT1 (R&D, 6514-SL-050), SPON1 (R&D, 3135-SP-025), DCN (R&D, 143-DE-100) at a concentration of 1ng/ul or LN511 (1 ug/cm^2^, Biolamina) for 4h at 37°C and cells grown on the coated plates between days 22 and 28.

### Immunocytochemistry, image acquisition and quantification

Cells were fixed in 4% PFA, washed in PBS and blocked in PBTA (PBS, 5% normal donkey serum (Jackson ImmunoResearch), 0.3% Triton X-100, and 1% BSA) for 1 hr at room temperature (RT). Primary antibodies were diluted PBTA and incubations were carried out overnight at 4°C. The primary antibodies used were the following: TH (1:500, Pel-Freez, P40101), Ki67 (1:1000, Cell Signaling Technology, 9449), NR4A2 (1:250, SantaCruz, sc990), bIII-tubulin (1:1000, Promega, G7121), LMX1(1:500, Millipore, AB10533), FOXA2(1:200, R&D, AF2400), Click-iT^TM^ EdU (Invitrogen, C10337) and Click-iT^TM^ Plus TUNEL (Invitrogen, C10617). Corresponding secondary antibodies were Alexa Fluor Dyes (Invitrogen) (1:1000) and incubated at RT for 2 hr. Cells were counterstained with DAPI (Thermo Fisher Scientific, D1306). Cells were then washed with PBS and stored at 4°C in Mounting medium (Dako).

Immunofluorescent images were captured with a confocal microscope (Zeiss LSM700 for hLT-NES experiments, Zeiss LSM980-Airy for hESC experiments) with a 20x 0.45 NA objective. A minimum of six images were per well were captured, with two technical replicates per condition in independent biological experiments. Quantifications of EdU and DAPI positive nuclei were performed using Qupath (Bankhead et al., 2017). Total TH cells as well as double positive TH/EdU and TH/DAPI cells were counted manually using Fiji (Schindelin et al., 2012). For each independent biological replicate, raw counts were min-max normalized to account for variations within each biological experiment.

### Statistical analysis

Results are presented as mean ± standard error of the mean (SEM). Significance of differences was assessed using Student’s t-tests, with Bonferroni-Holm multiple testing correction when applicable.

## AUTHOR CONTRIBUTIONS

ESA and EMT and performed and interpreted the biological validation experiments with support from CS. EMT, GLM, LFB, KL and ESA performed and interpreted the computational work. DG and CV, embryonic tissue dissections. PRdVC, ESA and CG, lentivirus work. CPP and ESA, cell culture counts. GLM and SI, RNA extraction, cDNA library preparation and NGS pipeline. CS, SL and GLM contributed to experimental design and interpretation of the results. ESA, EMT and EA designed experiments, interpreted results and wrote the manuscript. All authors have read and approved the manuscript.

## ACKNOWLEDGMENTS

We thank members of the Arenas lab for help, suggestions, and helpful discussions. Knut and Alice Wallenberg Foundation for support to the CLICK imaging facility at KI. Financial support was obtained from the Swedish Research Council (VR projects: DBRM, 2008:2811, 2011-3116 and 2011-3318, 2016-01526, 2020-01426), Swedish Foundation for Strategic Research (SRL program and SB16-0065), European Commission (NeuroStemCellRepair (602278), DDPD-Genes, Neurostemcell-Reconstruct (874758), PreciseCell PD (ERC-ADG 884608)), Knut and Alice Wallenberg Foundation (2018.0232), Karolinska Institutet (StratRegen SFO 2018) and Hjärnfonden (FO2019-0068) and Cancerfonden (CAN 2016/572), to EA. Chan Zuckerberg Initiative and the Silicon Valley (NDCN 2018-191929) to EA and SL. Grants from the Swedish Research Council and DDPDGENES to S.L and a fellowship from the Swedish Research Council to E.M.T.

## Supplementary figure legends

**Figure S1.**
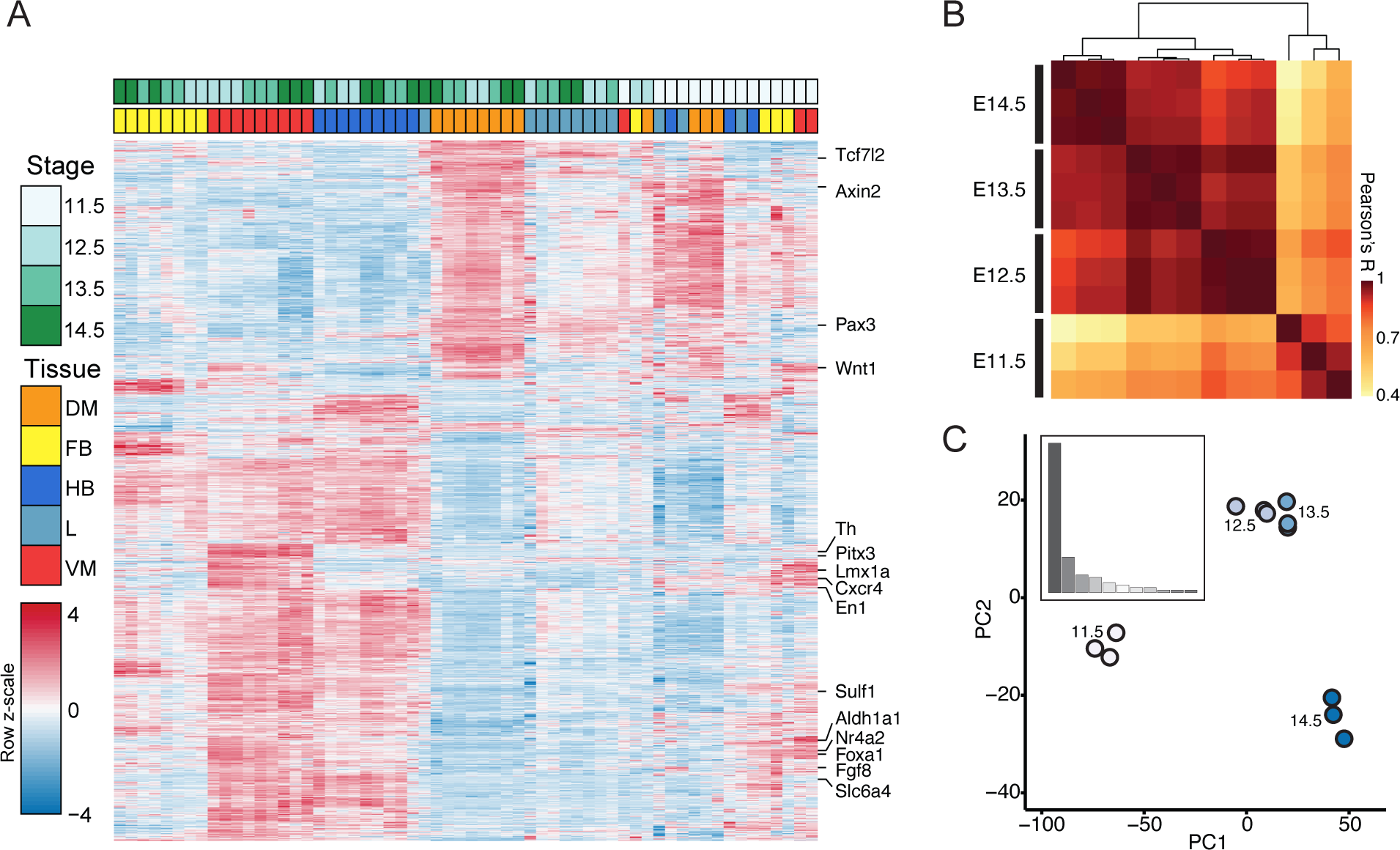
Bulk RNA-seq analysis of the mouse VM and neighboring brain regions during development. **(A)** Heat map representation of high variance genes (rows) in all the samples (columns) in different tissues in our dataset, from embryonic day 11.5 to 14.5. Samples are color coded by stage and region. **(B)** Pearson’s correlation of VM samples after filtering for variance (12.5%). **(C)** Principal component analysis of VM samples at different developmental times. Insert shows the percentage of variance by component.

**Figure S2.**
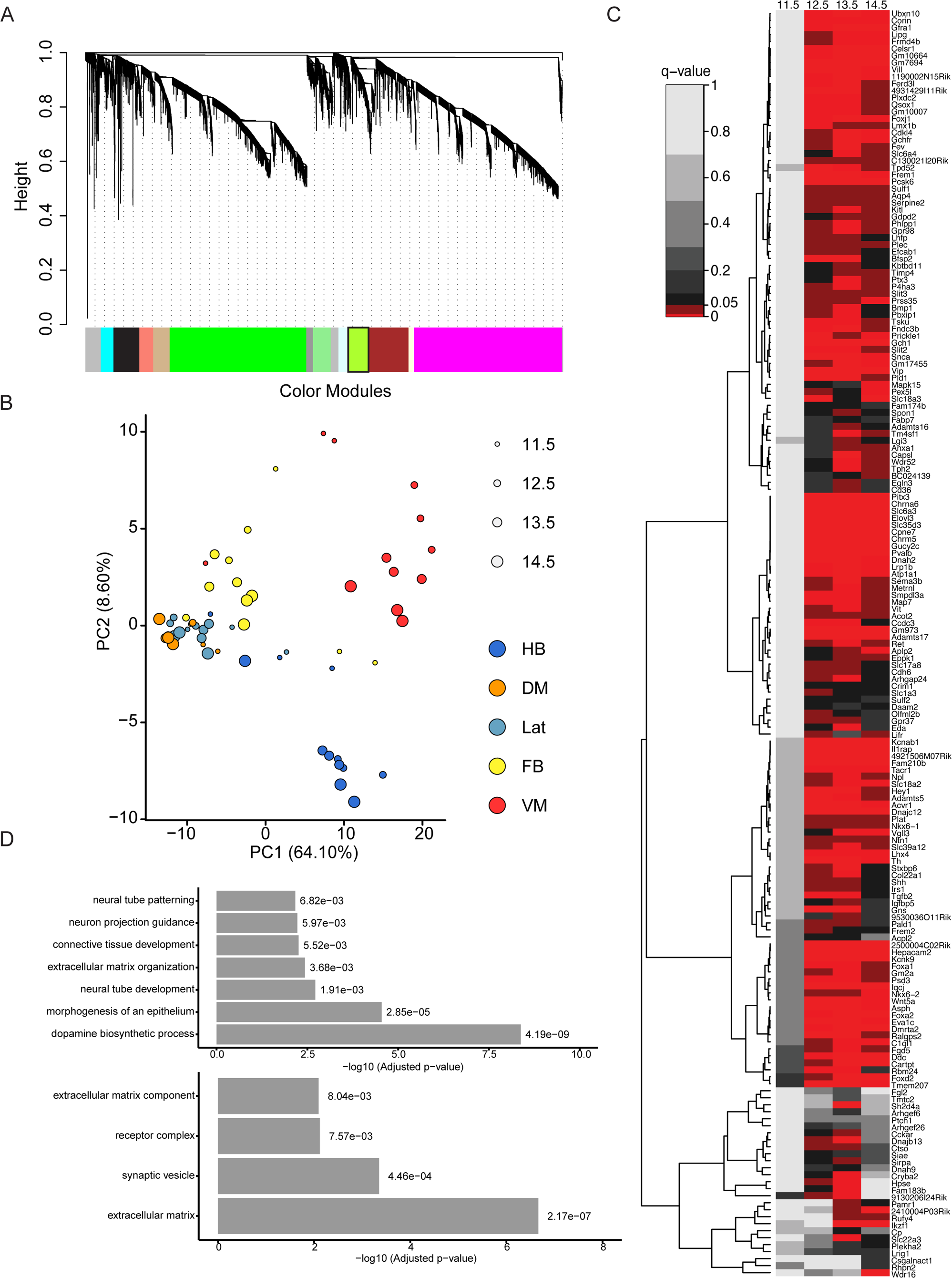
Weighted Gene Co-Expression Network Analysis of all bulk RNA-seq samples. **(A)** Dendrogram of modules obtained from weighted gene co-expression network analysis. mDA module is shown in light green and has a black border. **(B)** Principal component analysis of all the tissues and stages analyzed. The genes in the mDA module “light green” were sufficient to separate the VM from the rest of the tissue samples. Color represents tissue and size represents developmental stage. **(C)** Heat map of DEGs in the mDA module for each developmental stage. Red scale, q-values between 0 and 0.05; black to light gray scale, q-values above 0.05. **(D)** GO analysis of genes in the mDA module: Top, biological processes; Bottom, cellular components.

**Figure S3.**
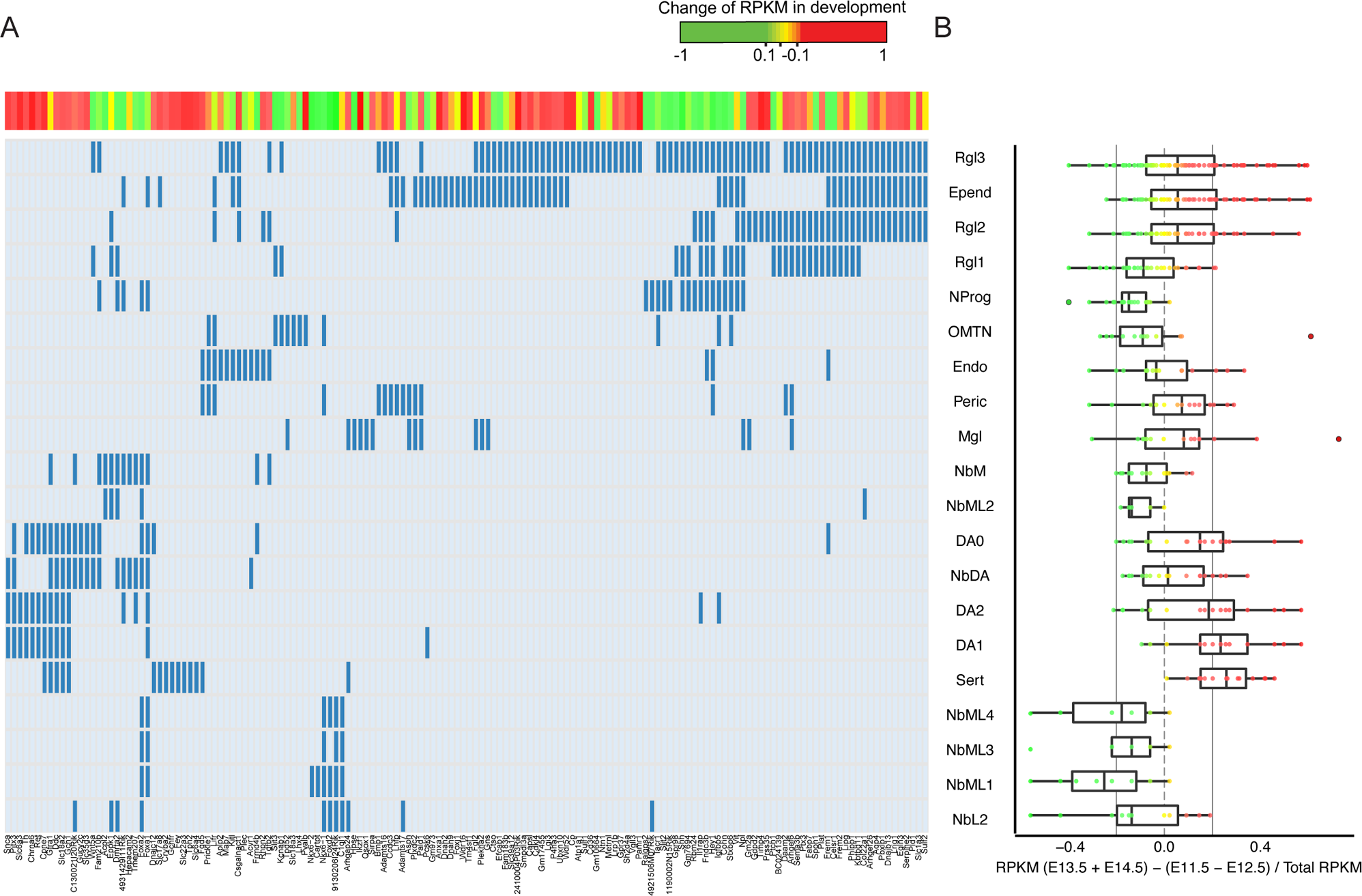
Single-cell network deconvolution. **(A)** Identity matrix of VM cell types and genes expressed in the mDA module. Blue lines represent significantly expressed genes in the corresponding cell type, compared to baseline. Red, gene increased; Yellow, maintained; Green, decreased over time. Color scale corresponds to figure 1B. **(B)** Analysis of gene expression levels (RPKM) in the mDA module for each cell type, comparing early (E11.5+E12.5) versus late stages (E13.5+E14.5).

**Figure S4.**
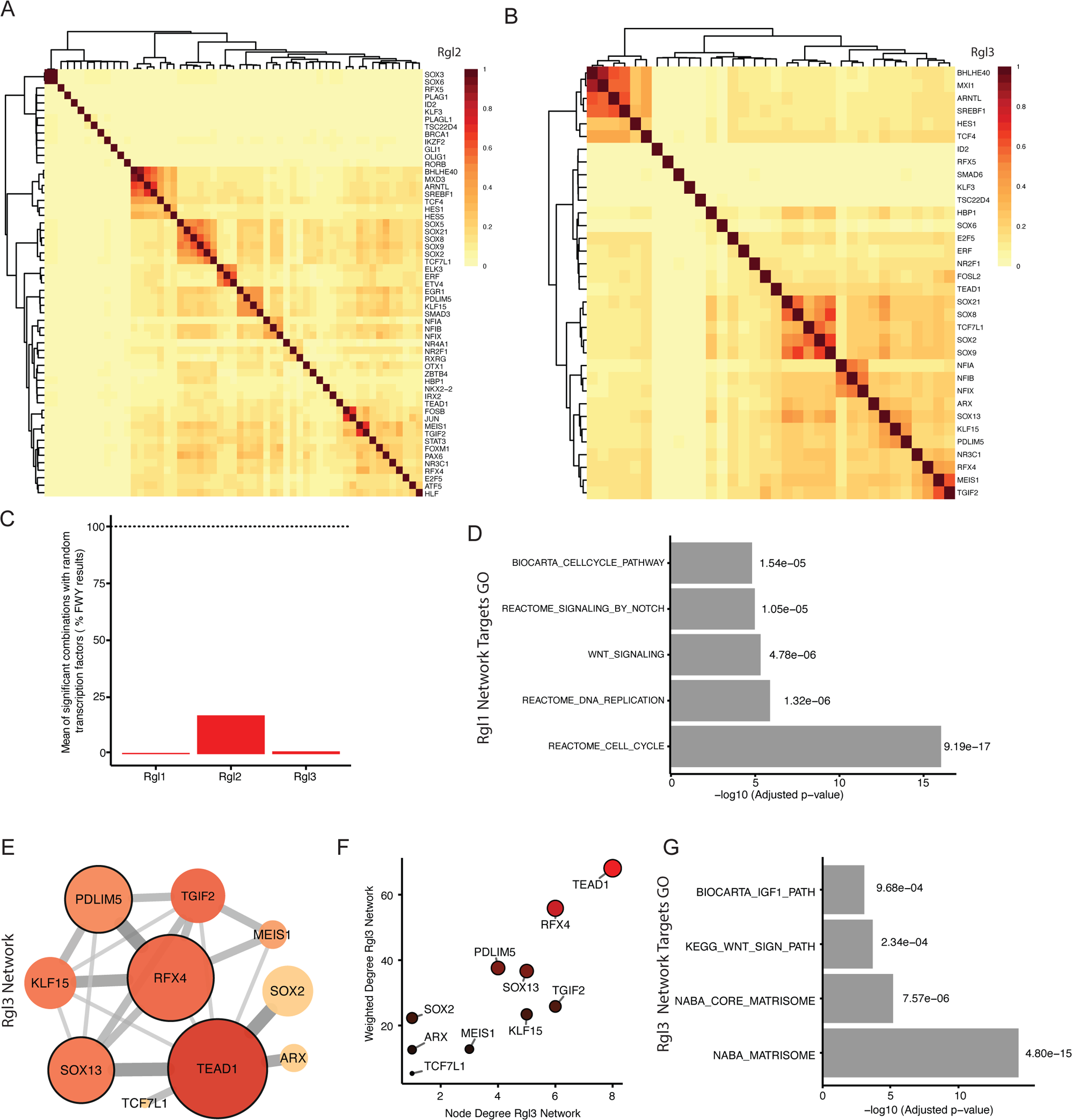
Combinatorial enrichment of Rgl2 and Rgl3 transcription factors. (**A, B**) Clustering of transcription factors expressed in Rgl2 (A) or Rgl3 (B) by Jaccard index of shared target genes. **(C)** Percentage of significant FWY combinations obtained with random selection of transcription factors while maintaining Rgl1-3 transcriptome constant. **(D)** GO terms for target genes of core transcription factors in Rgl1 TF network. Analysis was done with MSigDB gene set C2 canonical pathways v5.0. **(E)** Network representation of FWY analysis of Rgl3. Node color and size are proportional to node degree. Nodes with higher weighted degrees (core nodes) have a black border. Color intensity and width of the lines connecting the nodes are proportional to the interaction score for each transcription factor pair. **(F)** Plot comparing node degree and weighted node degree of the network obtained by FWY analysis of Rgl3. Color intensity and size are proportional to the weighted degree. **(G)** GO terms for target genes of core transcription factors in Rgl3 TF network. Analysis was done with MSigDB gene set C2 canonical pathways v5.0.

**Figure S5.**
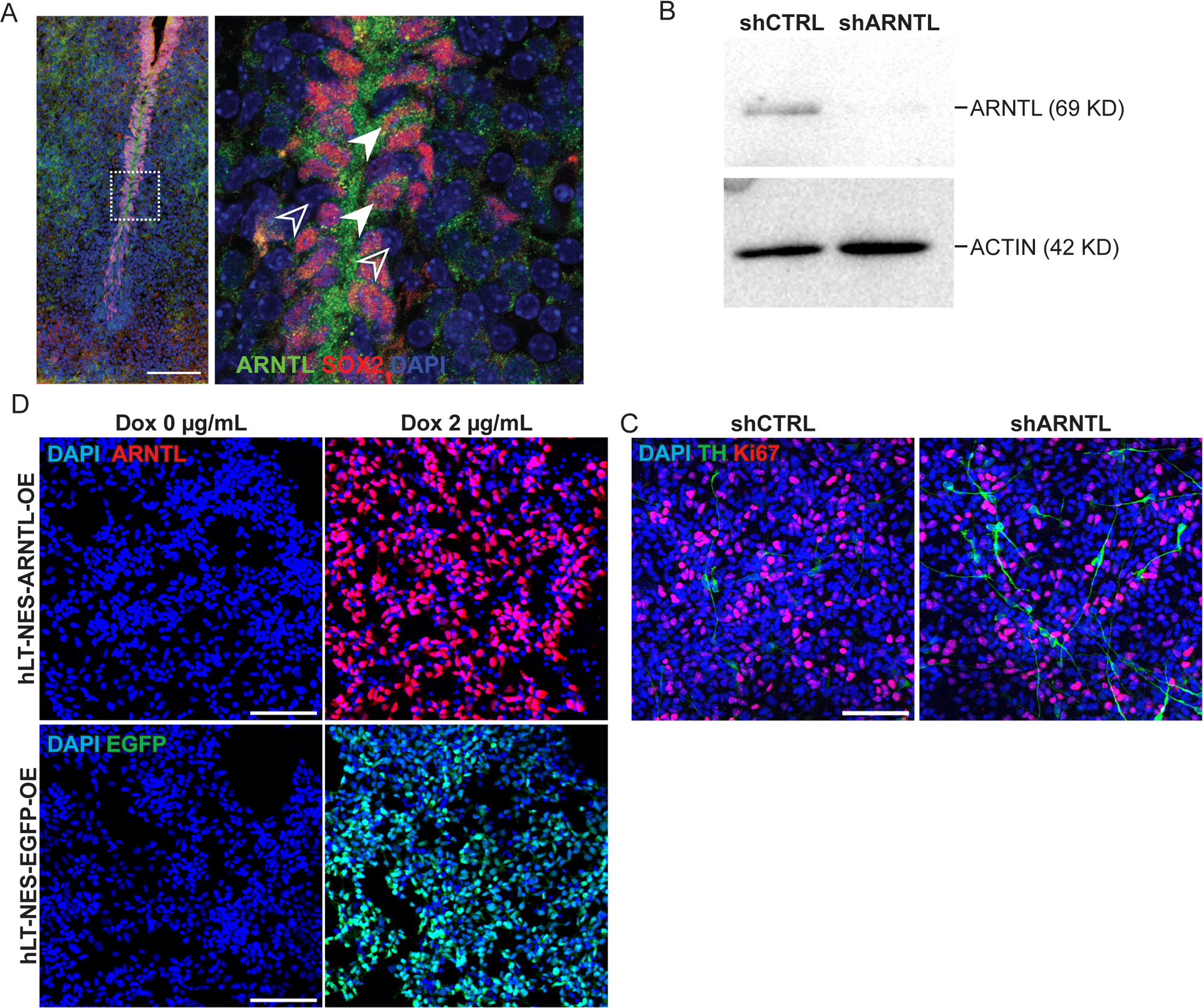
Validation of ARNTL knockdown and over-expression in hLT-NES cells. **(A)** ARNTL protein (green) is present in SOX2+ cells (red) in the developing mouse VM at E13.5. DAPI in blue. Image to the right magnified from white box on the left. Full arrow heads, double positive SOX2 and ARNTL cells. Empty arrowheads, SOX2 and ARNTL negative cells. Scale bar, 100 µm. **(B)** Western blot analysis identified the presence of ARNTL in control hLT-NES cells. ARNTL was dramatically reduced by shRNAs against ARNTL (upper blot, second lane). ACTIN was used as loading control (lower blot). **(C)** Immunostaining for TH^+^ neurons and Ki67^+^ proliferating progenitor cells in shCTRL and shARNTL hLT-NES on day 4 of the differentiation protocol. Scale bar, 100 µm. **(D)** ARNTL or EGFP transgene expression was induced by doxycycline treatment (2 µg/mL) for 24 hours in stable hLT-NES ARNTL or EGFP inducible cell lines, respectively. Scale bar, 100 µm.

**Figure S6.**
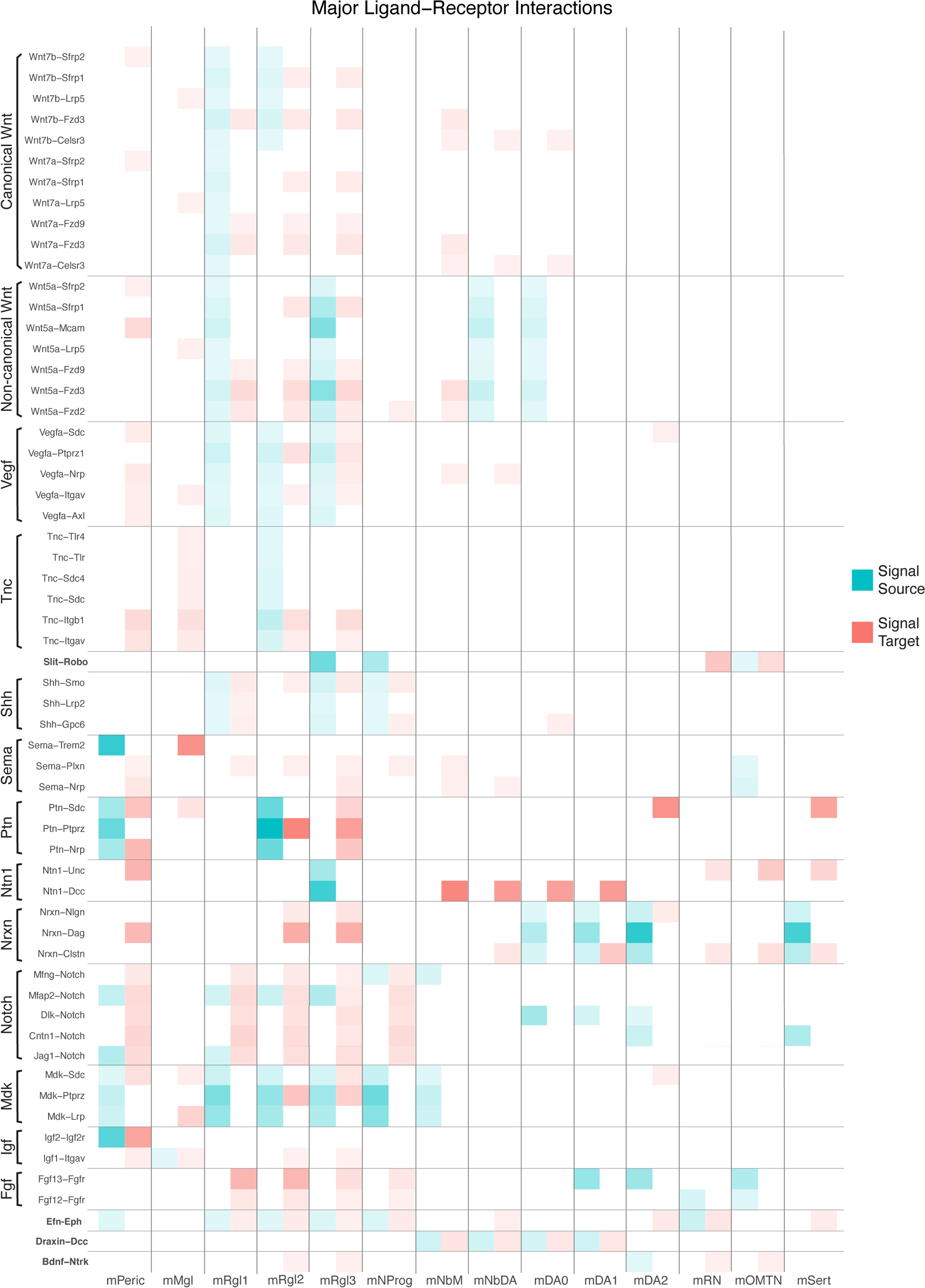
Overview of outgoing and incoming signaling interactions in the developing mouse VM. (A) Heatmap of major predicted ligand-receptor interactions in cell types of the developing VM filtered by Bayesian posterior probability of gene expression. Only cells with predicted interactions are shown. Pathways are indicated to the left. Interactions involving ligands or receptors of the Notch, Fgf, Slit, Robo, Nrxn, Ntn1, Efn-Eph, Sema, Mdk, Ptn, Tnc, Vegfa families are shown together (e.g. Notch: Notch1-4; Dlk: Dlk1-2; Fgfr: Fgfr1-3; Slit: Slit1-3; Robo: Robo1, 2, 4; Nrxn: Nrxn1-3; Nxph1/Nlgn1; Unc: Unc5a-d; Efn: Efna1,2,5, Efnb1,2; Eph: Epha1-5, Ephb1-3; Nrp: Nrp1,2; Sema: Sema3c,d,f,g/4c,g/5a/6a,b,c/7a; Plxn: Plxna/b1-4; Mdk-Sdc2,4; Mdk-Lrp1; Ptn-Sdc2,3,4; Ptn-Nrp1; Tnc-Sdc1,4; Vegfa-Nrp1; Vegfa-Sdc2,3,4).

**Figure S7.**
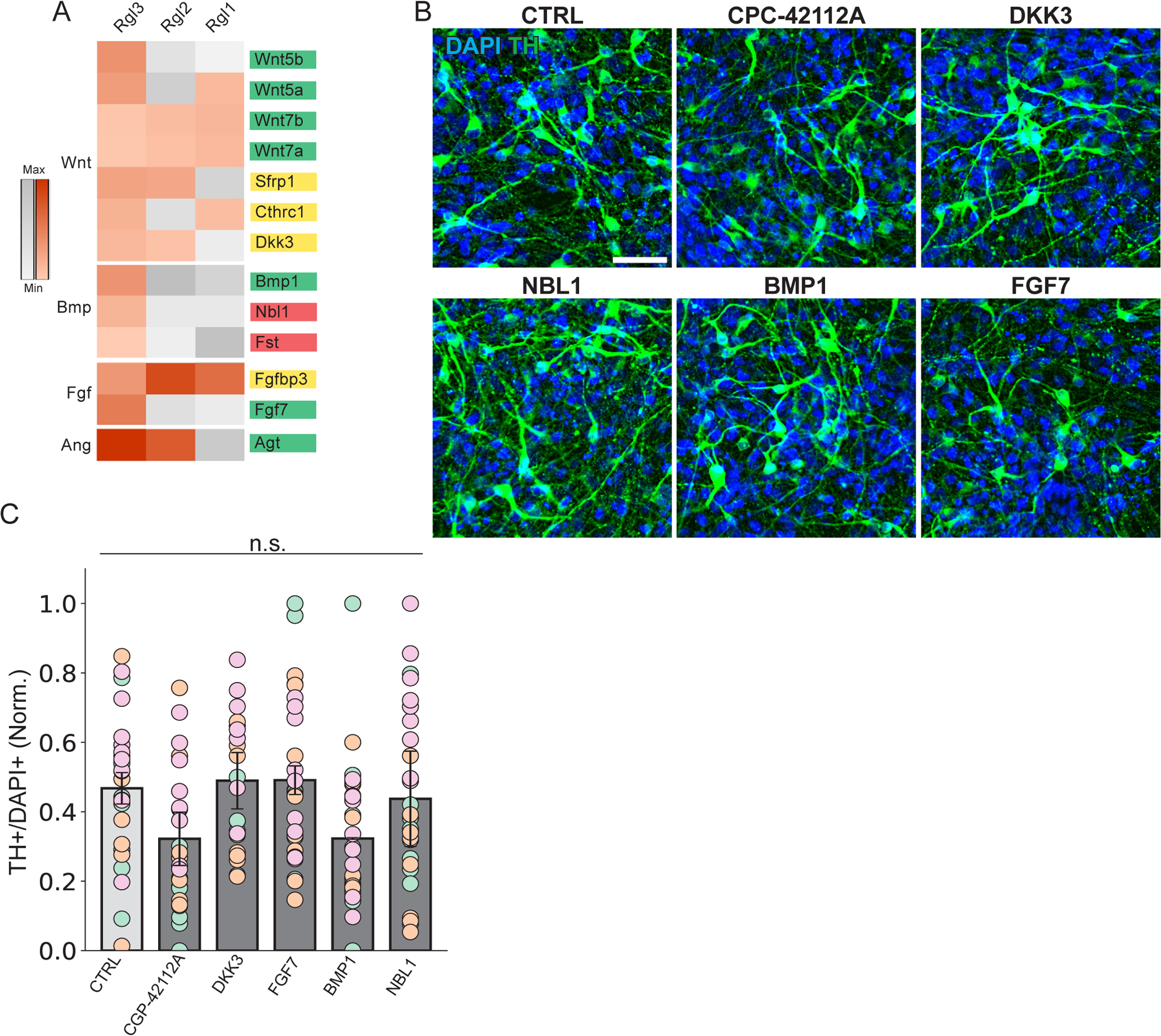
Influence of selected Rgl3 ligands on TH neurons derived from hESCs. **(A)** Heat map representation of selected ligands expressed by Rgl cell types in the developing VM. Ligands are colored according to their activity; green for activators, red for inhibitors and yellow for context-dependent modulation. Pathway is indicated on the left. Color intensity is proportional to the Bayesian estimate of expression level. Gray scale indicates values below significance level. **(B)** Immunostaining of TH^+^ neurons derived from hESCs that were cultured in the presence of the selected ligands/small molecules from days 22-35 and analyzed on day 35 of differentiation. Scale bar, 100 µm. **(C)** Quantification of TH^+^/DAPI cells on day 35. Data is normalized to the average of each experiment. n.s., non-significant. Multiple testing correction was performed using Bonferroni-Holm correction. Each data point is represented by a circle, and the circle’s color corresponds to the biological replicate (n = 3 independent experiments).

**Figure S8.**
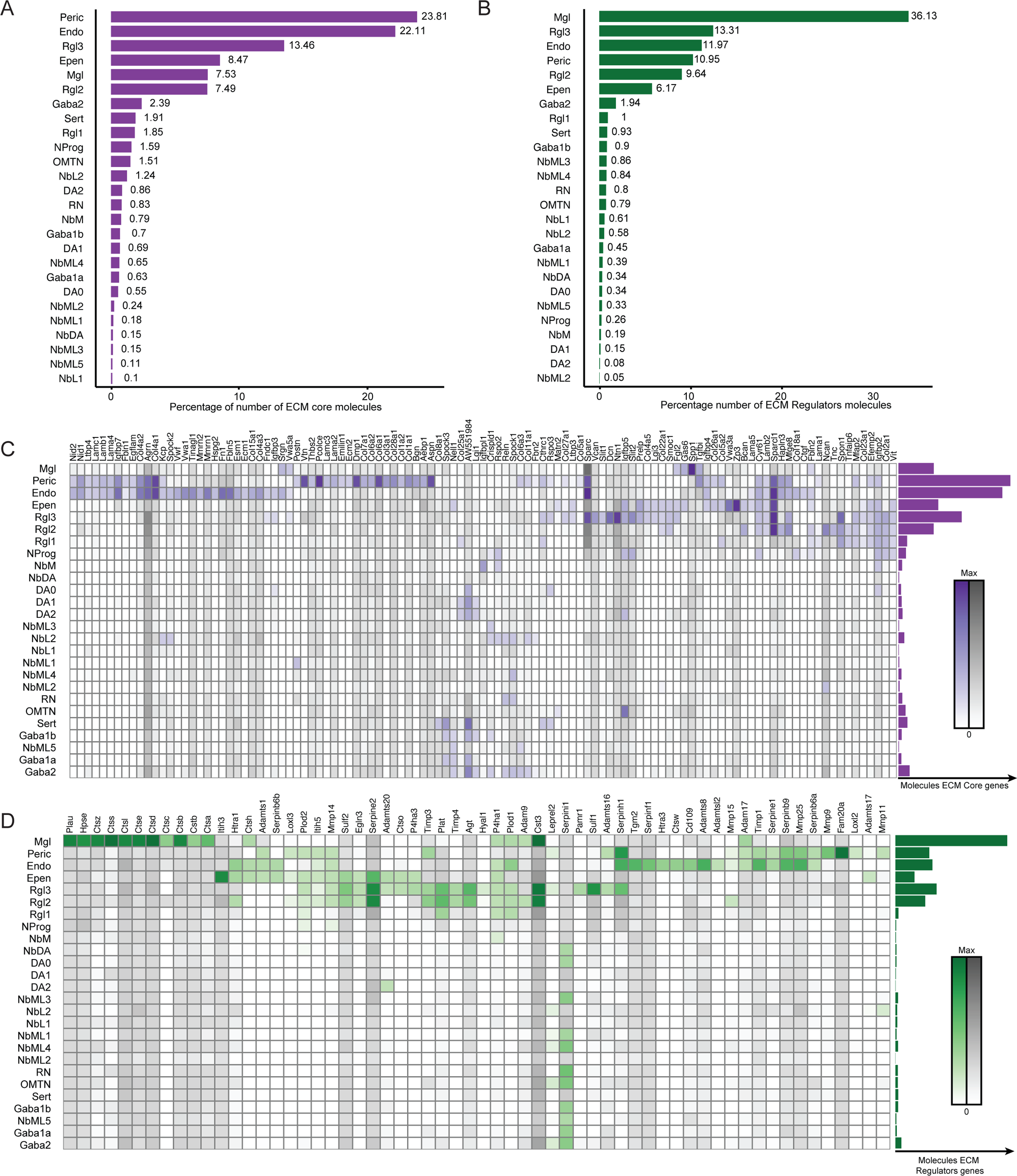
Contribution of individual cell types to ECM core components and ECM regulation during mouse VM development. **(A, B)** Bar plot showing the contribution of each VM cell type to the total number of transcripts for core ECM genes (**A**) and regulatory ECM genes **(B)** between days 11.5-18.5 of mouse VM development. **(C, D)** Heat maps showing the expression levels of ECM regulators **(C)** and ECM core components **(D)** in all VM cell types. Color intensity is proportional to the Bayesian estimate of expression level. Gray scale indicates values below significance level. Bar plot to the right represents total average of transcripts for ECM core components/regulators per cell type.

**Figure S9.**
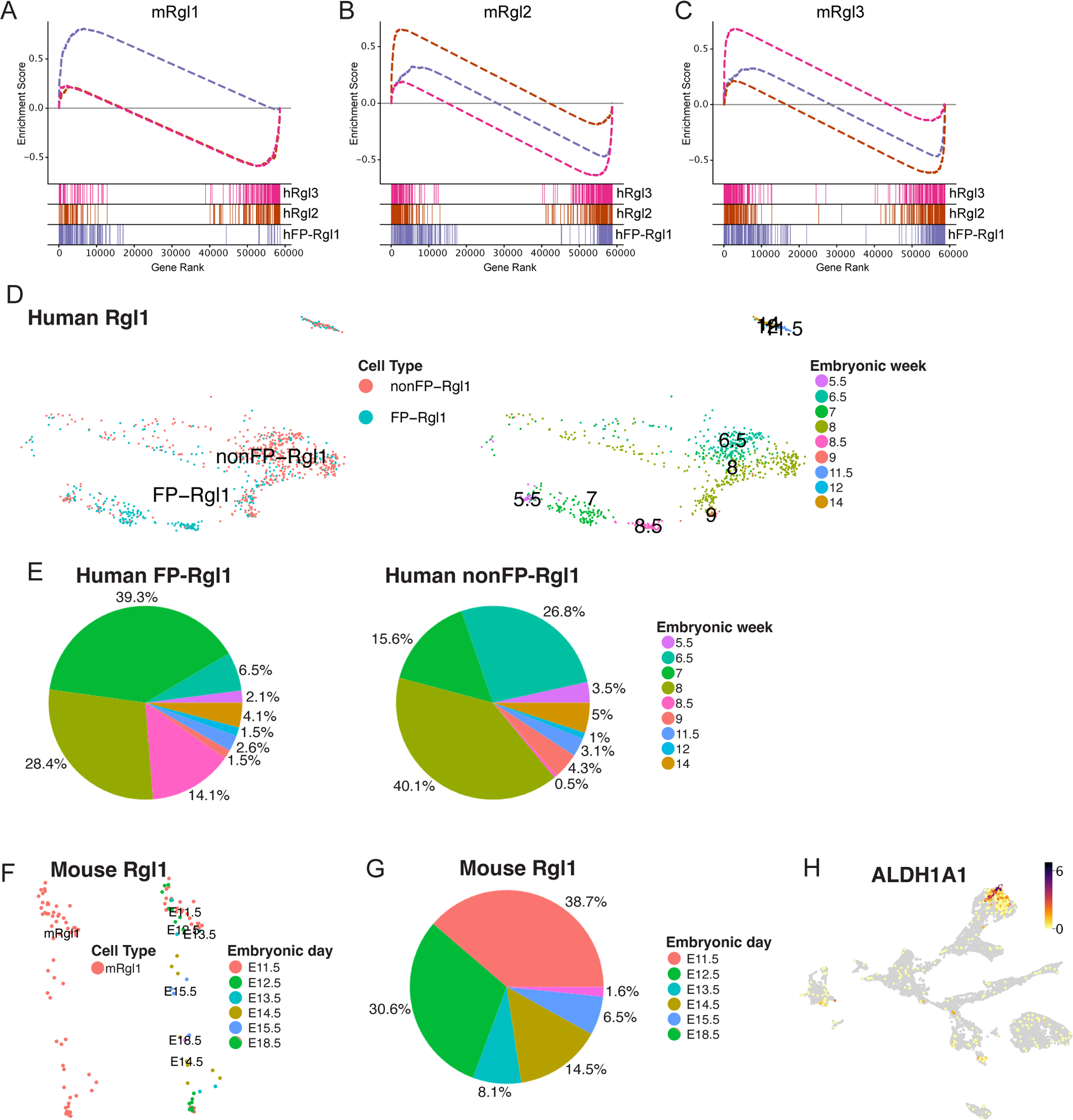
Comparison of human and mouse Rgl cells used for CellChat, and definition of the *ALDH1A1+* population in the developing human VM. **(A-C)** Gene set enrichment analysis of human FP-Rgl1, hRgl2 and hRgl3 transcriptomes compared to genes unique in mouse Rgl1 (**A**), mRgl2 (**B**) and mRgl3 (**C**). **(A)** hRgl1: NES = 1.369, FDR < 0.001. hRgl2: NES = −1.115, FDR < 0.001. hRgl3: NES = −1.115, FDR < 0.001 **(B)** hRgl1: NES = −1.171, FDR = 0.172. hRgl2: NES = 1.524, FDR < 0.001. hRgl3: −1.300, FDR = 0.035. **(C)** hRgl1: NES = −1.159, FDR = 0.171. hRgl2: NES = −1.330, FDR = 0.018. hRgl3: NES = 1.578, FDR < 0.001. **(D)** UMAP of human FP-Rgl1 and nonFP-Rgl1 cells used for CellChat analysis colored by embryonic week. **(E)** Pie chart showing the embryonic week of human FP-Rgl1 and nonFP-Rgl1 cells used for CellChat analysis. **(F)** UMAP of mouse Rgl1 cells used for CellChat analysis colored by embryonic day. **(G)** Pie chart showing the embryonic day of mouse Rgl1 cells used for CellChat analysis. **(H)** UMAP showing ALDH1A1 expression in the developing human VM using the data from Braun et al. 2023.

